# Multivalency transforms SARS-CoV-2 antibodies into broad and ultrapotent neutralizers

**DOI:** 10.1101/2020.10.15.341636

**Authors:** Edurne Rujas, Iga Kucharska, Yong Zi Tan, Samir Benlekbir, Hong Cui, Tiantian Zhao, Gregory A. Wasney, Patrick Budylowski, Furkan Guvenc, Jocelyn C. Newton, Taylor Sicard, Anthony Semesi, Krithika Muthuraman, Amy Nouanesengsy, Katherine Prieto, Stephanie A. Bueler, Sawsan Youssef, Sindy Liao-Chan, Jacob Glanville, Natasha Christie-Holmes, Samira Mubareka, Scott D. Gray-Owen, John L. Rubinstein, Bebhinn Treanor, Jean-Philippe Julien

## Abstract

The novel severe acute respiratory syndrome coronavirus 2 (SARS-CoV-2), which causes Coronavirus Disease 2019 (COVID-19), has caused a global pandemic. Antibodies are powerful biotherapeutics to fight viral infections; however, discovery of the most potent and broadly acting clones can be lengthy. Here, we used the human apoferritin protomer as a modular subunit to drive oligomerization of antibody fragments and transform antibodies targeting SARS-CoV-2 into exceptionally potent neutralizers. Using this platform, half-maximal inhibitory concentration (IC_50_) values as low as 9 × 10^−14^ M were achieved as a result of up to 10,000-fold potency enhancements. Combination of three different antibody specificities and the fragment crystallizable (Fc) domain on a single multivalent molecule conferred the ability to overcome viral sequence variability together with outstanding potency and Ig-like *in vivo* bioavailability. This MULTi-specific, multi-Affinity antiBODY (Multabody; or MB) platform contributes a new class of medical countermeasures against COVID-19 and an efficient approach to rapidly deploy potent and broadly-acting therapeutics against infectious diseases of global health importance.

**One Sentence Summary:** multimerization platform transforms antibodies emerging from discovery screens into potent neutralizers that can overcome SARS-CoV-2 sequence diversity.

## Introduction

The continuous threat to public health from respiratory viruses such as the novel SARS-CoV-2 underscores the urgent need to rapidly develop and deploy prophylactic and therapeutic interventions to combat pandemics. Monoclonal antibodies (mAbs) have been used effectively for the treatment of infectious diseases as exemplified by palivizumab for the prevention of respiratory syncytial virus in high-risk infants^1^ or Zmapp, mAb114 and REGN-EB3 for the treatment of Ebola^2^. Consequently, mAbs targeting the Spike (S) protein of SARS-CoV-2 have been a focus for the development of medical countermeasures against COVID-19. To date, several antibodies targeting the S protein have been identified^3–19^ and several are under clinical evaluation^20,21^. Receptor binding domain (RBD)-directed mAbs that interfere with binding to angiotensin converting enzyme 2 (ACE2), the receptor for cell entry^22^, are usually associated with high neutralization potencies^11,12,14^.

mAbs can be isolated by B-cell sorting from infected donors, immunized animals or by identifying binders in pre-assembled libraries. Despite these methodologies being robust and reliable for the discovery of virus-specific mAbs, identification of the best antibody clone can be a lengthy process. For example, RNA viruses have a high mutation rate. Indeed, 198 naturally occurring mutations in the S protein of SARS-CoV-2 have been already registered in the GISAID database^23^. Identification of broadly neutralizing mAbs (bnAbs) that target the most conserved epitopes to overcome viral escape mutations is therefore critical. This approach has been employed to discover bnAbs against human immune deficiency virus 1 (HIV-1)^24^, Influenza^25^, and Ebola^26^. However, their identification required extensive sampling and high-throughput sequencing and consequently several years of research. Hence, there is an unmet need for the development of a platform that bridges antibody discovery and the rapid identification and deployment of highly potent and broadly neutralizing mAbs.

The potency of an antibody is greatly affected by its ability to simultaneously interact multiple times with its epitope^27–29^. This enhanced apparent affinity, known as avidity, has been previously reported to increase the neutralization potency of nanobodies^30,31^ and IgGs over Fabs^9,16,18^ against SARS-CoV-2. In order to rapidly propel mAbs that emerge during initial screening efforts into potent neutralizers against SARS-CoV-2, we have developed an antibody-scaffold technology, using the human apoferritin protomer as a modular subunit, to multimerize antibody fragments. We demonstrate the ability of this technology to combine one to three different Fab specificities together with Fc on the same molecule. The resulting MB molecules can increase mAb potency by up to four orders of magnitude and resist sequence variability in the spike protein. Therefore, the MB offers a versatile “plug-and-play” platform to enhance anti-viral characteristics of prophylactic/therapeutic molecules in the fight against the SARS-CoV-2 pandemic, and to be quickly deployed in the setting of emerging pandemics.

## Results

### Avidity enhances neutralization potency

We used the self-assembly of the light chain of human apoferritin to multimerize antigen binding moieties targeting the SARS-CoV-2 S glycoprotein. Apoferritin protomers self-assemble into an octahedrally symmetric structure with an ~6 nm hydrodynamic radius (R_h_) composed of 24 identical polypeptides^32^. The N terminus of each apoferritin subunit points outwards of the spherical nanocage and is therefore accessible for the genetic fusion of proteins of interest. Upon folding, apoferritin protomers act as building blocks that drive the multimerization of the 24 proteins fused to their N termini (**Fig. 1a**).

**Fig. 1.**
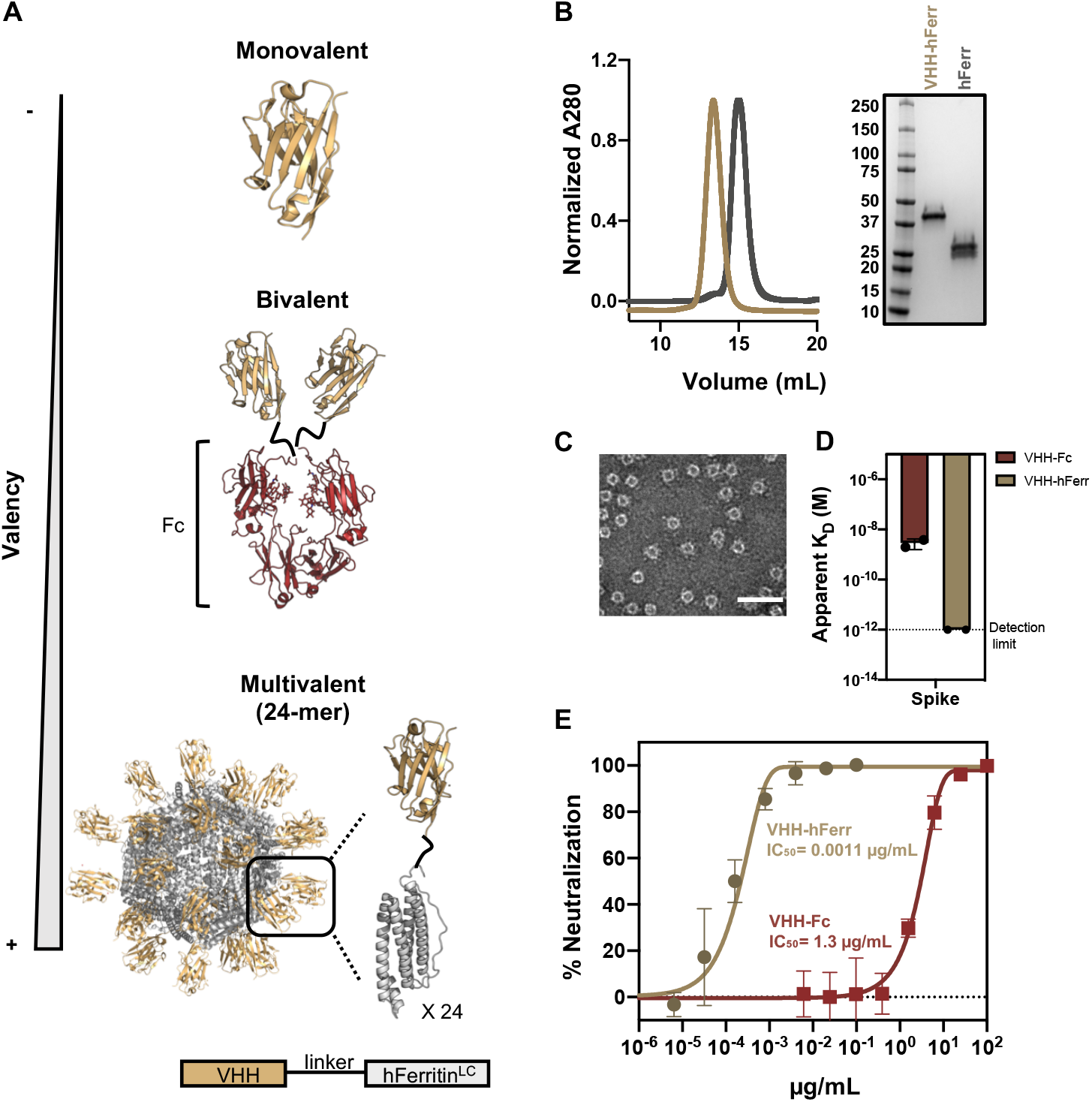
Avidity enhances binding and neutralization of VHH against SARS-CoV-2. (**A**) Schematic representation of a monomeric VHH domain and its multimerization using a conventional Fc scaffold or human apoferritin. (**B**) Size exclusion chromatography and SDS-PAGE of apoferritin alone (*gray*) and VHH-72 apoferritin particles (*gold*). (**C**) Negative stain electron microscopy of VHH-72 apoferritin particles. (Scale bar 50 nm). (**D**) Comparison of the binding avidity (apparent K_D_) of VHH-72 to SARS-CoV-2 S protein when displayed in a bivalent (red) or 24-mer (gold) format. Apparent K_D_ higher than 10^−12^ M (dash line) is beyond the instrument detection limit. (**E**) Neutralization potency against SARS-CoV-2 PsV (color coding is as in d). One representative out of two experiments with similar results is shown.

First, we investigated the impact of multivalency on the ability of the single chain variable domain VHH-72 to block viral infection. VHH-72 has been previously described to neutralize SARS-CoV-2 when fused to a Fc domain, but not in its monovalent format^30^. The light chain of human apoferritin displaying 24 copies of VHH-72 assembled into monodisperse, well-formed spherical particles (**Fig. 1b-c**) and showed an enhanced binding avidity to the S protein **(Fig. 1d)** in comparison to bivalent VHH-72-Fc. Strikingly, display of VHH-72 on the light chain of human apoferritin achieved a 1000-fold increase in neutralization potency against SARS-CoV-2 pseudovirus (PsV) compared to the conventional Fc fusion (**Fig. 1e**).

### Multabodies have IgG-like properties

The Fc confers *in vivo* half-life and effector functions to IgGs through interaction with neonatal Fc receptor (FcRn) and Fc gamma receptors (FcγR), respectively. To confer these IgG-like properties to the MB, we next sought to incorporate both Fabs and Fc domains. Because a Fab is a hetero-dimer consisting of a light and a heavy chain, and the Fc is a homodimer, we created single-chain Fab (scFab) and Fc (scFc) polypeptide constructs to allow for their direct fusion to the N terminus of the apoferritin protomer. As a first step, we generated a species-matched surrogate molecule that consists of mouse light chain apoferritin fusions to a mouse scFab and a mouse scFc (IgG2a subtype), or a modified mouse scFc (LALAP mutation) to assess their, biodistribution, immunogenicity and pharmacokinetics *in vivo* (**Fig. S1**). As expected, binding affinity of the WT MB to mouse Fc receptors was enhanced in comparison to the parental IgG and the LALAP mutation reduced binding affinity to FcγR1 (**Fig. S1a**) Subcutaneous administration of these MBs in C57BL/6 or BALB/c mice was well tolerated with no decrease in body weight or visible adverse events. *In vivo* bioavailability and biodistribution of the MB were similar to the parental IgG (**Fig. S1b and c**). Confirming the role of the Fc in mediating *in vivo* bioavailability, serum half-life could be extended by mutations in the Fc (**Fig. S1b**). Presumably because all sequences derived from the host, the MB did not induce an anti-drug antibody response in mice **(Fig. S1d).**

In view of these results, we aimed to generate MBs derived from the previously reported IgG BD23^5^ and 4A8^6^ that target the SARS-CoV-2 spike RBD and N-terminal domain (NTD), respectively. Addition of scFcs into the MB reduces the number of scFabs that can be multimerized. In order to endow the MB platform with Fc without compromising Fab avidity and hence neutralization potency, we engineered the apoferritin protomer to accommodate more than 24 components per particle. Based on its four-helical bundle fold, the human apoferritin protomer was split into two halves: the two N-terminal α helices (N-Ferritin) and the two C-terminal α helices (C-Ferritin). In this configuration, the scFc fragment of human IgG1 and the scFab of anti-SARS-CoV-2 IgGs were genetically fused at the N terminus of each apoferritin half, respectively. Split apoferritin complementation led to hetero-dimerization of the two halves and in turn, full apoferritin self-assembly displaying more than 24 copies of scFc and scFab on the nanocage periphery (**Fig. 2a**). Conveniently, this design allows for the straightforward purification of the MB using Protein A akin to IgG purification.

**Fig. 2.**
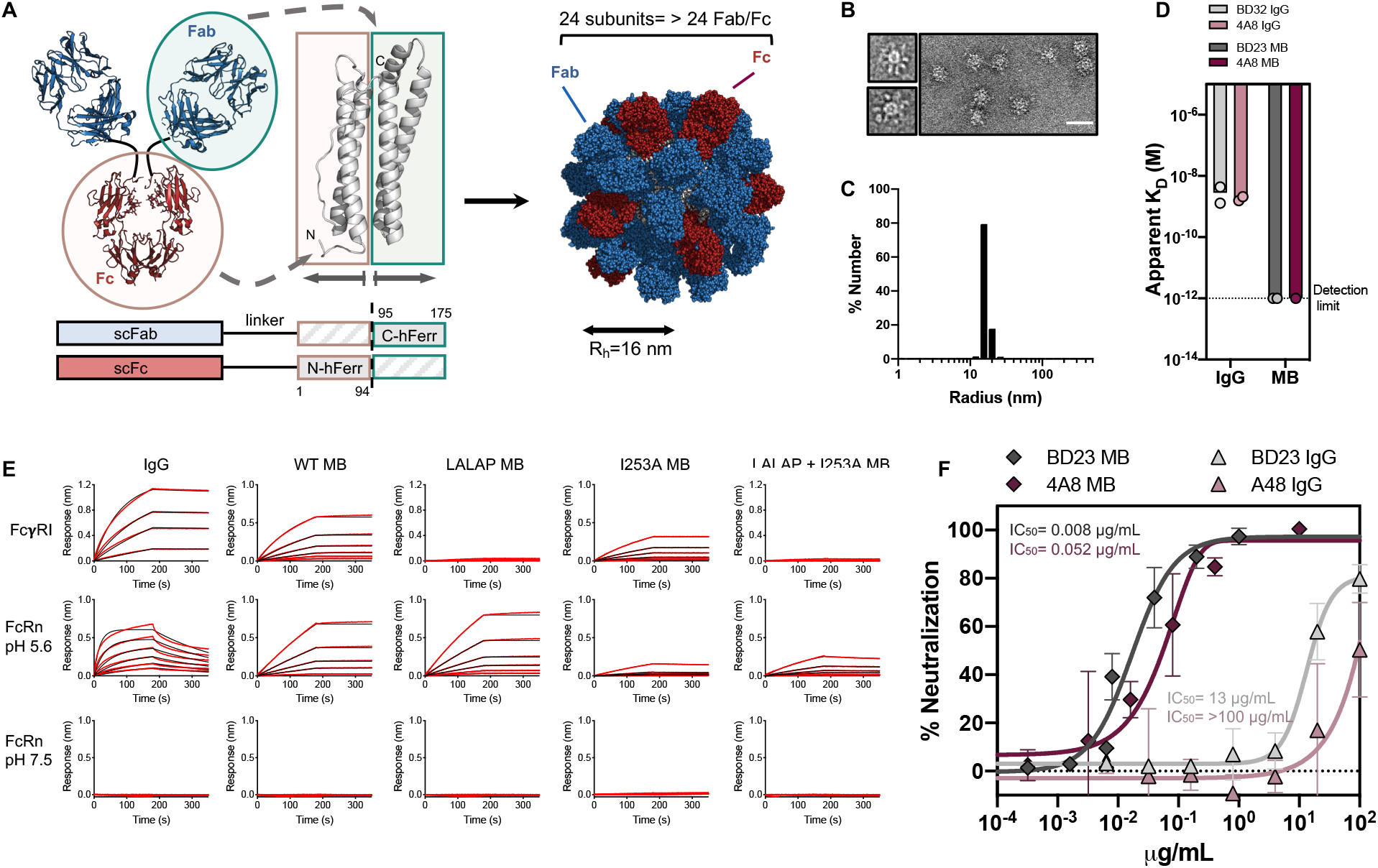
Protein engineering to multimerize IgG-like particles against SARS-CoV-2. (**A**) Schematic representation of the human apoferritin split design. (**B**) Negative stain electron micrograph of the MB. (Scale bar 50 nm). (**C**) Hydrodynamic radius (R_h_) of the MB. (**D**) Avidity effect on the binding (apparent K_D_) of A48 (*purple*) and BD23 (*gray*) to the SARS-CoV-2 Spike. (**E**) Kinetic curves of BD23 IgG and MB with different Fc sequence variants binding to FcγRI (top row), FcRn at endosomal pH (middle row) and FcRn at physiological pH (bottom row). Black lines represent raw data whereas red lines represent global fits. (**F**) Neutralization of SARS-CoV-2 PsV by A48 and BD23 IgGs and MBs. The mean values ± SD for two technical replicates for a representative experiment is shown.

This split MB design forms 16 nm hydrodynamic radius (R_h_) spherical particles with an uninterrupted ring of density and regularly spaced protruding scFabs and scFc (**Fig. 2b and 2c**). Hence, the MB is on the lower size range of natural IgMs^33^ but packs more weight on a similar size to achieve multi-valency and multi-specificity. Binding kinetics experiments demonstrated that avidity of the MB for the Spike was preserved upon addition of Fc fragments **(Fig. 2d).** Binding to human FcγRI and FcRn at endosomal pH confirmed that scFc was properly folded in the split MB design and that LALAP and I253A mutations lowered binding affinities to FcγRI and FcRn, respectively (**Fig. 2e**). SARS-CoV-2 PsV neutralization assays with the split design MBs showed that enhanced binding affinity for the S protein translates into an improved neutralization potency in comparison to their IgG counterparts, with a ~1600-fold and ~2500-fold increase for BD23 and A48, respectively **(Fig. 2f)**. Combined, this data supported the further exploration of the MB as an IgG-like platform that confers extraordinary multivalency.

### From antibody discovery to ultrapotent neutralizers

We next assessed the ability of the MB platform to transform mAb binders identified from initial phage display screens into potent neutralizers against SARS-CoV-2 **(Fig. 3a)**. Following standard biopanning protocols, 20 human mAb binders with moderate affinities that range from 10^−6^ to 10^−8^ M against the RBD of SARS-CoV-2 were selected **(Table S1)**. These mAbs were produced as full-length IgGs and MBs and their capacity to block viral infection was compared in a neutralization assay against SARS-CoV-2 PsV **(Fig. 3b** and **Fig. S2a)**. Notably, MBs preserved the thermostability of their parental IgGs (**Fig. S2b-c**) and significantly enhanced the potency of 18 out of 20 (90%) IgGs by up to four orders of magnitude. The largest increment was observed for mAb 298 which went from an IC_50_ of ~30 μg/mL as an IgG to 0.0001 μg/mL as a MB. Strikingly, 11 mAbs were converted from non-neutralizing IgGs to neutralizing MBs in the tested concentration ranges. Seven MBs displayed exceptional potency with IC_50_ values between 0.2-2 ng/mL against SARS-CoV-2 PsV using two different target cells (293T-ACE2 and HeLa-ACE2 cells) (**Fig. 3b** and **Fig. S3a**). The enhanced neutralization potency of the MB was further confirmed with authentic SARS-CoV-2 virus for the mAbs with the highest potency (**Fig. 3c** and **Fig. S3b**). Retrospectively, all IgGs and MBs were tested for their ability to bind to the S protein and the RBD of SARS-CoV-2 **(Fig. S4)**. Increased avidity resulted in higher apparent binding affinities with no detectable off-rates against the S protein, most likely due to inter-spike crosslinking that translates into high neutralization potency (**Fig. 3b-d** and **Fig. S4**). Overall, the data shows that the MB platform is compatible with rapid delivery of ultrapotent IgG-like molecules.

**Fig. 3.**
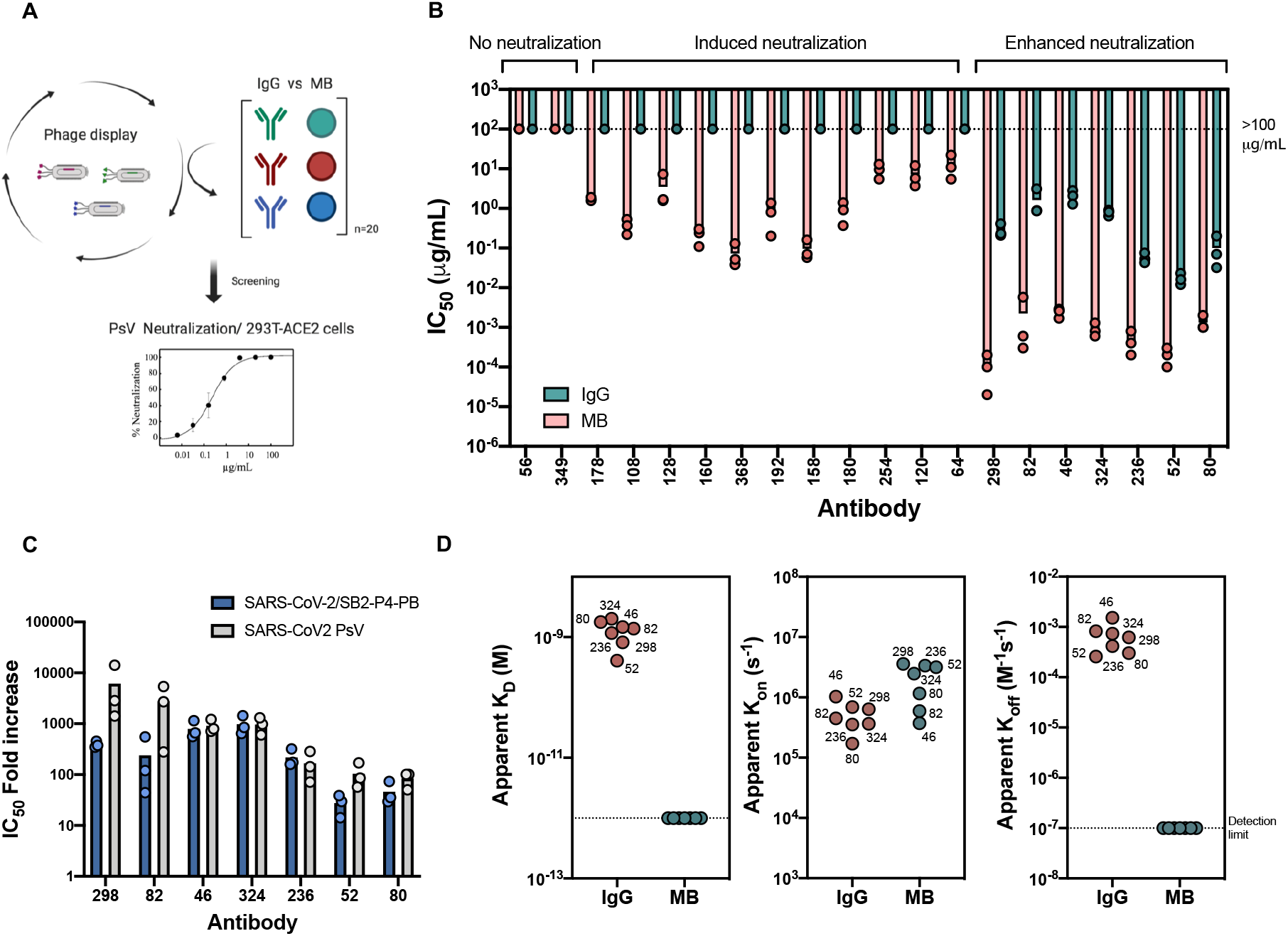
The Multabody enhances the potency of human mAbs from phage display. (**A**) Work flow for the identification of potent anti-SARS-CoV-2 neutralizers using the MB technology. (**B**) Comparison of neutralization potency between IgGs and MBs that display the same human Fab sequences derived from phage display. (**C**) IC_50_ values fold increase upon multimerization. (**D**) Apparent affinity (K_D_), association (k_on_) and dissociation (k_off_) rates of the most potent neutralizing MBs compared to their IgG counterparts for binding the SARS-CoV-2 S protein. Three biological replicates and their mean are shown for IC_50_ values in (b) and (c).

### Epitope mapping

Based on their neutralization potency, seven mAbs were selected for further characterization: 298 (IGHV1-46/IGKV4-1), 82 (IGHV1-46/IGKV1-39), 46 (IGHV3-23/IGKV1-39), 324 (IGHV1-69/IGKV1-39), 236 (IGHV1-69/IGKV2-28), 52 (IGHV1-69/IGKV1-39) and 80 (IGHV1-69/IGKV4-1) (**Fig. 3b** and **Table S1)**. Epitope binning experiments showed that these mAbs target two main sites on the RBD, with one of these bins overlapping with the ACE2 binding site **(Fig. 4a** and **Fig. S5)**. Cryo-EM structures of Fab-SARS-CoV-2 S complexes at a global resolution of ~6-7 Å confirmed that mAbs 324, 298, 80 bind overlapping epitopes **(Fig. 4b, Fig. S6a-c** and **Table S2**). To gain insight into the binding of mAbs targeting the second bin, we obtained the cryo-EM structure of Fab 46 in complex with the RBD at a global resolution of 4.0 Å **(Fig. 4c, Fig. S6d** and **Table S2**), and the crystal structure of Fabs 298 and 52 as a ternary complex with the RBD at 2.95 Å resolution (**Fig. 4d, Fig. S7** and **Table S3**).

**Fig. 4.**
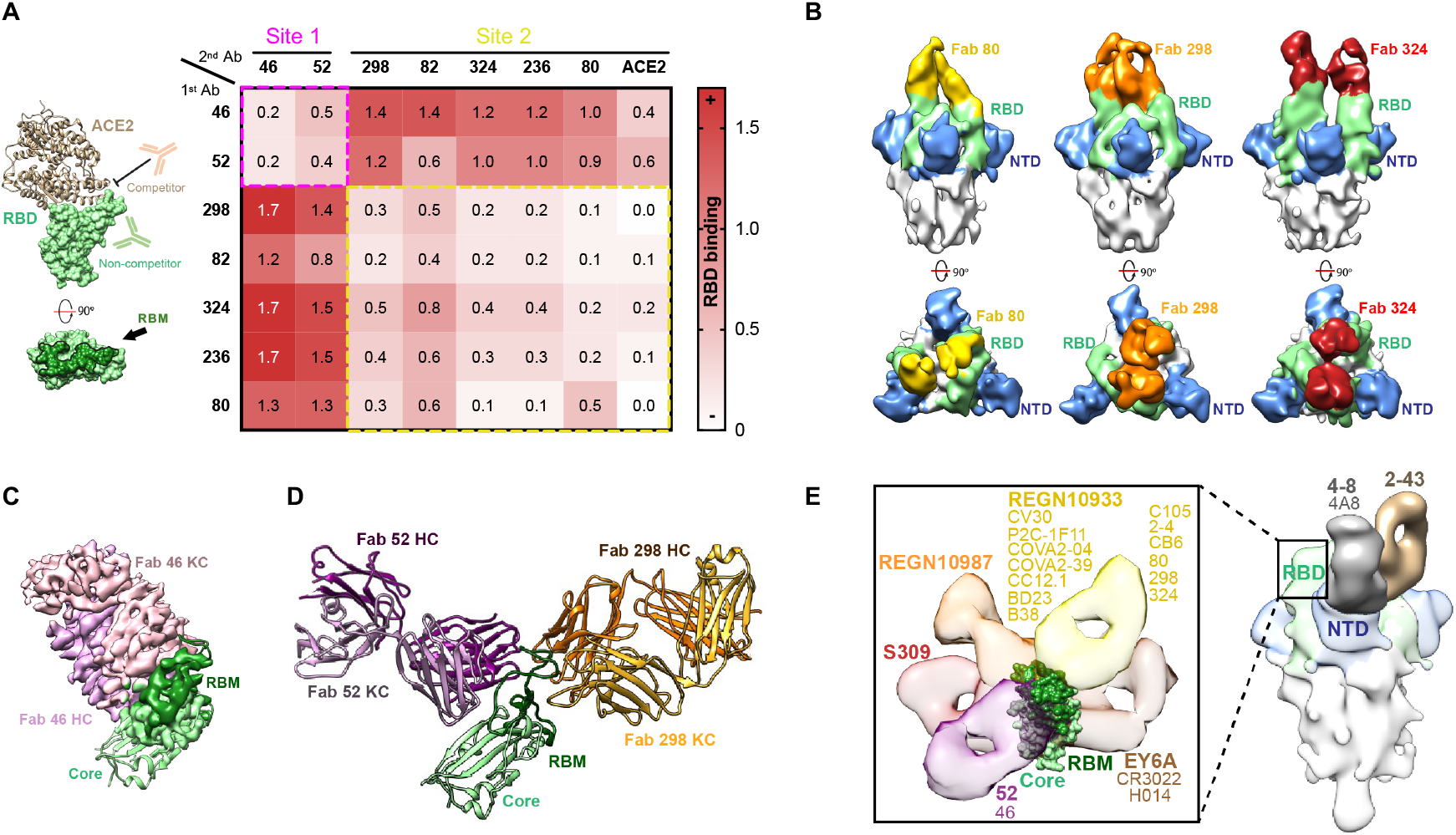
Epitope delineation of the most potent mAb specificities. (**A**) Surface and cartoon representation of RBD (*light green* for the core and *dark green* for RBM) and ACE2^46^ (*light brown*) binding. Heat map showing binding competition experiments. Epitope bins are highlighted by dashed-line boxes. (**B**) 15.0 Å filtered cryo-EM reconstruction of the Spike (*grey*) in complex with Fab 80 (*yellow*), 298 (*orange*) and 324 (*red*). The RBD and NTD are shown in *green* and *blue*, respectively. (**C**) Cryo-EM reconstruction of Fab 46 (*pink*) and RBD (*green*) complex. A RBD cartoon^46^ is fitted into the partial density observed for the RBD. (**D**) Crystal structure of the ternary complex formed by Fab 52 (*purple*), Fab 298 (*orange*) and RBD (*green*). (**E**) Composite image depicting antibodies targeting SARS-CoV-2 S with available PDB or EMD entries^3,4,6,8,10,16,17,47,48^. Inset: close up view of antibodies targeting different antigenic sites on the RBD. The mAb with the lowest reported IC_50_ value against SARS-CoV-2 PsV was selected as a representative antibody of the bin (color coding as in b).

The crystal structure shows that mAb 298 binds almost exclusively to the ACE2 receptor binding motif (RBM) of the RBD (residues 438-506). In fact, out of 16 RBD residues involved in binding mAb 298, 12 are also involved in ACE2-RBD binding **(Fig. S7a-c** and **Table S4 and S5**). The RBM is stabilized by 11 hydrogen bonds from heavy and light chain residues of mAb 298. In addition, RBM Phe486 is contacted by 11 mAb 298 residues burying approximately 170 Å^2^ (24% of the total buried surface area on RBD) and hence is central to the antibody-antigen interaction **(Fig. S7a** and **Table S5**).

Detailed analysis of the RBD-52 Fab interface reveals that the epitope of mAb 52 is shifted towards the core of the RBD encompassing 20 residues of the RBM and 7 residues in the core domain **(Fig. 4c, Fig. S7b** and **Table S6** and **S7**). In agreement with the competition data, antibody 52 and antibody 46 share a similar binding site, although they approach the RBD with slightly different angles **(Fig. 4c-d** and **Fig. S7d**). Inspection of previously reported structures of RBD-antibody complexes reveal that antibodies 46 and 52 target a site of vulnerability on the SARS-CoV-2 spike that has not been described previously **(Fig. 4e)**. The epitope targeted by these antibodies is partially occluded by the NTD in the S “closed” conformation, suggesting that the mechanism of action for this new class of antibodies could involve Spike destabilization. Together, these data demonstrate that the enhanced potency observed for the MB platform is associated with mAbs that target both overlapping and distinct epitope bins.

### Multabodies overcome Spike sequence variability

To explore whether MBs could resist viral escape, we tested the effect of four naturally occurring RBD mutations (L452R, A475V, N439K and V483A)^34^ **(Fig. 5a)** on the binding and neutralization of the seven human mAbs of highest potency **(Fig. 5b-c)**. In addition, the impact of mutating Asn234, a glycosylation site, to Gln was also assessed because the absence of glycosylation at this site has been previously reported to decrease sensitivity to neutralizing antibodies targeting the RBD^34^. The more infectious PsV variant D614G^35^ was also included in the panel. Mutation L452R significantly decreased binding and potency of antibodies 52 and 46, while antibody 298 was sensitive to mutation A475V **(Fig. 5b-c)**. Deletion of the N-linked glycan at position Asn234 increased viral resistance to the majority of the antibodies, especially mAbs 46, 80 and 324, emphasizing the important role of glycans in viral antigenicity **(Fig. 5c)**. Strikingly, the following mAbs were minimally impacted in their exceptional neutralization potency by any S mutation when present in the MB format: 298, 80, 324, and 236 **(Fig. 5d)**. Mutation L452R decreased the sensitivity of the 46-MB and 52-MB but in contrast to their parental IgGs, they remained neutralizing against this PsV variant **(Fig. 5d)**. The more infectious SARS-CoV-2 PsV variant D614G was neutralized with similar potency as the WT PsV for both IgGs and MBs **(Fig. 5c** and **Fig. S8a**).

**Fig. 5.**
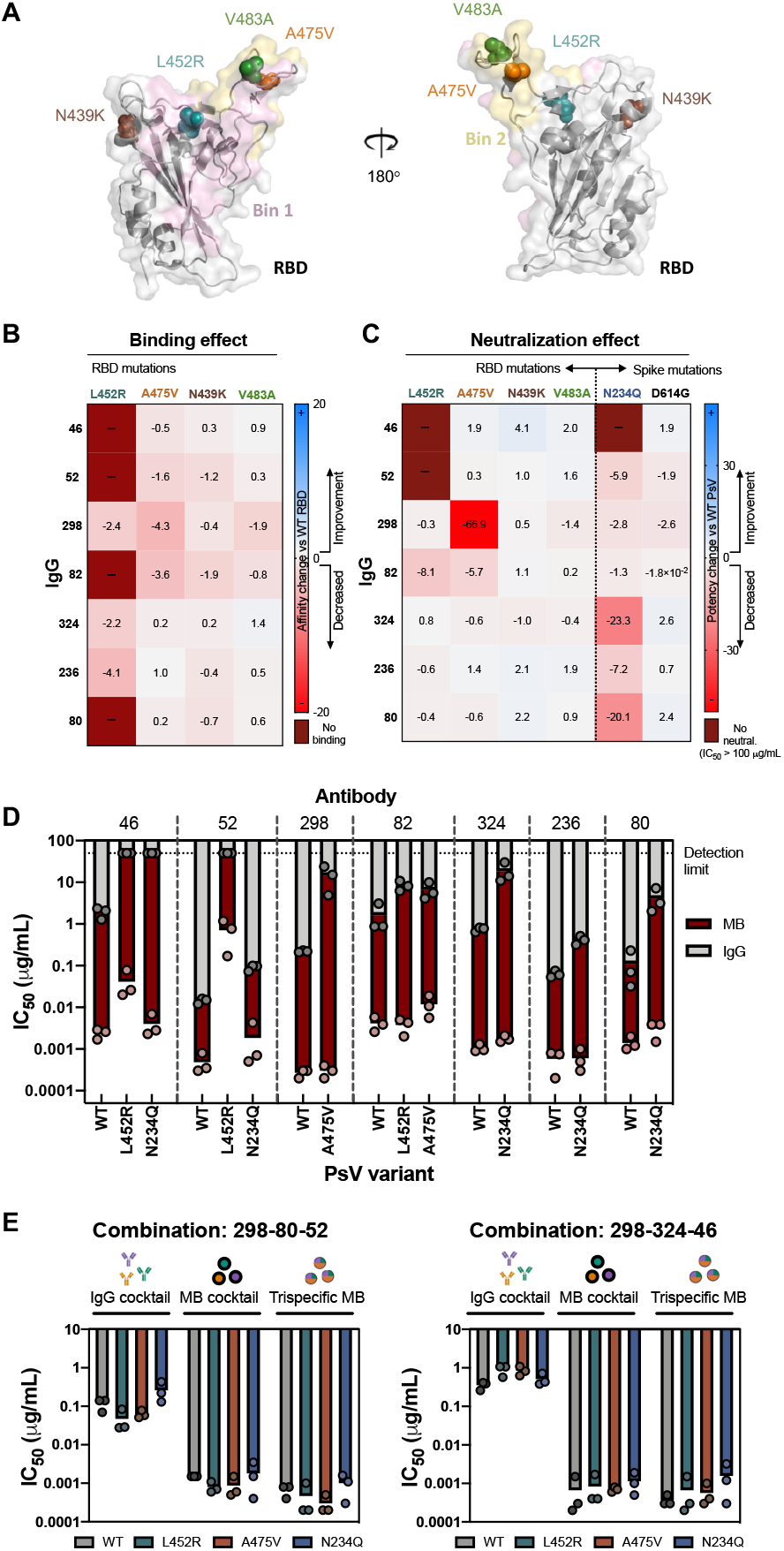
Multabodies overcome SARS-CoV-2 sequence diversity. (**A**) Cartoon representation of the RBD showing four naturally occurring mutations as spheres. The epitopes of mAbs 52 (*light pink*) and 298 (*yellow*) are shown as representative epitopes of each bin. (**B**) Affinity and (**C**) IC_50_ fold-change comparison between WT and mutated RBD and PsV, respectively. (**D**) Neutralization potency of IgG (*grey bars*) vs MB (*dark red bars*) against SARS-CoV-2 PsV variants in comparison to WT PsV. (**E**) Neutralization potency comparison of two IgG cocktails (three IgGs), monospecific MB cocktails (three MBs) and tri-specific MBs against WT SARS-CoV-2 PsV and variants. mAbs sensitive to one or more PsV variants were selected to generate the cocktails and the tri-specific MBs. The mean of three biological replicates are shown in (b), (c), (d) and (e).

MB cocktails consisting of three monospecific MB resulted in pan-neutralization across all PsV variants without a significant loss in potency and hence achieved a 100-1000-fold higher potency compared to the corresponding IgG cocktails **(Fig. 5e** and **Fig. S8c-d**). In order to achieve breadth within a single molecule, we next generated tri-specific MBs by combining multimerization subunits displaying three different Fabs in the same MB assembly **(Fig. S8b)**. Notably, the resulting tri-specific MBs exhibited pan-neutralization while preserving the exceptional neutralization potency of the monospecific versions **(Fig. 5e** and **Fig. S8c-d**). The most notable increase in potency was for the 298-324-46 combination, where the IgG cocktail neutralized WT SARS-CoV-2 PsV with an ~0.5 μg/mL (3.3 × 10^−9^ M) IC_50_, while the tri-specific MB achieved broad neutralization with an exceptional IC_50_ of ~0.0005 μg/mL (2.2 × 10^−13^ M). The potency enhancement of MB cocktails and tri-specific MBs in comparison to their corresponding IgG tri-specific cocktails was also observed with live replicating SARS-CoV-2 virus **(Fig. S8e**).

## Discussion

We have developed an antibody-multimerization platform to increase avidity of mAbs targeting SARS-CoV-2. The seven most potent MBs have IC_50_ values of 0.2 to 2 ng/mL (9 × 10^−14^ to 9 × 10^−13^ M) against SARS-CoV-2 PsVs and therefore are, to our knowledge, within the most potent antibody-like molecules reported to date against SARS-CoV-2.

The MB platform was designed to include key favorable attributes from a developability perspective. First, the ability to augment antibody potency is independent of antibody sequence, format or epitope targeted. The modularity and flexibility of the platform was exemplified by enhancing the potency of a VHH and multiple Fabs that target non-overlapping regions on two SARS-CoV-2 S sub-domains (RBD and NTD). Second, in contrast to other approaches to enhance avidity such as tandem fusions of single chain variable fragments^36,37^, which suffer from low stability, MBs self-assemble into highly stable particles with aggregation temperatures values similar to those of their parental IgGs. Third, alternative multimerization strategies like streptavidin^38^, verotoxin B subunit scaffolds^39^ or viral-like nanoparticles^40^ face immunogenicity challenges and/or poor bioavailability because of the absence of an Fc fragment and therefore the inability to undergo FcRn–mediated recycling. The light chain of apoferritin is fully human, biologically inactive, has been engineered to include Fc domains, and despite multimerization of >24 Fab/Fc fragments, has a hydrodynamic radius similar to an IgM. As such, a surrogate mouse MB did not elicit anti-drug antibodies in mice, was detectable in the sera for over a week, and has similar biodistribution as its parental IgG.

Different increases in neutralization potency were observed for different mAb sequences tested on the MB against SARS-CoV-2. This suggests that the ability of the MB to enhance potency may depend on epitope location on the Spike or the geometry of how the Fabs engage the antigen to achieve neutralization. The fact that 2 out of 20 SARS-CoV-2 RBD binders were not rescued by the MB platform also suggests limitations based on mAb sequences and binding properties. Nevertheless, the capacity of the MB to transform avidity into neutralization potency across a range of epitope specificities on the SARS-CoV-2 Spike highlights the potential for using this technology broadly. It will be interesting to explore the potency-enhancement capacity of the MB platform against viruses with low surface spike density like HIV^41^, or against other targets like the tumor necrosis factor receptor superfamily, where bivalency of conventional antibodies limits their efficient activation^42^.

Virus escape can arise in response to selective pressure from treatments or during natural selection. A conventional approach to combat escape mutants is the use of antibody cocktails targeting different epitopes. MBs showed a lower susceptibility to S mutations in comparison to their parental IgGs, presumably because the loss in affinity was compensated by enhanced binding avidity. Hence, when used in cocktails, the MB overcame viral sequence variability with exceptional potency. In addition, the split MB design allows combination of multiple antibody specificities within a single multimerized molecule resulting in similar potency and breadth as the MB cocktails. To our knowledge, these MBs represent the first tri-specifics described against SARS-CoV-2. Multi-specificity within the same particle could offer additional advantages such as intra-S avidity and synergy for the right combination of mAbs, setting the stage for further investigation of different combinations of mAb specificities on the MB.

Overall, the MB platform provides a tool to surpass antibody affinity limits and generate broad and potent neutralizing molecules without the need for extensive antibody discovery or engineering efforts. This platform will accelerate the timeline from discovery to development of antibodies as medical countermeasures against COVID-19 and in future pandemics.

## Acknowledgements

We thank F Krammer for providing the WT SARS-CoV-2 Spike plasmid. We thank JD Bloom and AC Gingras for access to 293T-ACE2 cells and reagents to make the SARS-CoV-2 PsV. We thank DR Burton for providing HeLa-ACE2 cells and the SARS-CoV-2 Spike mutant D614G. We are thankful to C Pettus, J Wang, D Pineda and K Patel for their work panning the SuperHuman 2.0 library against the SARS-CoV-2 RBD and characterizing binders. We thank JM Jorgensen and S Popa for assistance with the Octet RED96 and Unit Instruments. The following reagents were produced under HHSN272201400008C and obtained through BEI Resources, NIAID, NIH: Vector pCAGGS Containing the SARS-Related Coronavirus 2, Wuhan-Hu-1 Spike Glycoprotein Receptor Binding Domain (RBD), NR-52309, Vector pCAGGS Containing the SARS-Related Coronavirus 2, Wuhan-Hu-1 Spike Glycoprotein Gene (soluble, stabilized), NR-52394.

## Author contributions

ER, BT and JPJ conceived the research and designed the experiments; ER, IK, YZT, SB, HC, TZ, GW, PB, FG, JN, TS, AS, KM, AN, KP, SB, SY and SLC performed experimental work. JG, NCH, SM and SDGO provided critical reagents and expertise. ER, IK, YCT, BT, JLR and JPJ analyzed the data. ER and JPJ wrote the manuscript with input from all authors.

## Data and materials availability

The electron microscopy maps have been deposited at the Electron Microscopy Data Band (EMDB) with accession codes EMD-22738, EMD-22739, EMD-22740, EMD-22741 (Table S2). The crystal structure of 298-52-RBD complex (Table S3) is available from the Protein Data Bank under accession code PDB: 7K9Z. Sequences of the monoclonal antibodies are available in Table S1. Other data are available from the corresponding author upon reasonable request.

## Supplementary Materials

### Materials and Methods

#### Protein expression and purification

Genes encoding VHH-human apoferritin fusion, Fc fusions, Fabs, IgG, and RBD mutants were synthesized and cloned by GeneArt (Life Technologies) into the pcDNA3.4 expression vector. All constructs were expressed transiently in HEK293F cells (Thermo Fisher Scientific) at a density of 0.8 × 10^6^ cells/mL with 50 μg of DNA per 200 mL of cells using FectoPRO (Polyplus Transfections) in a 1:1 ratio unless specified otherwise. After 6-7 days of incubation at 125 rpm oscillation at 37° C, 8% CO_2_, and 70% humidity in a Multitron Pro shaker (Infors HT), cell suspensions were harvested by centrifugation at 5000 ×g for 15 min and supernatants were filtered through a 0.22 μm Steritop filter (EMD Millipore). Fabs and IgGs were transiently expressed by co-transfecting 90 μg of the LC and the HC in a 1:2 ratio and purified using KappaSelect affinity column (GE Healthcare) and HiTrap Protein A HP column (GE Healthcare), respectively with 100 mM glycine pH 2.2 as the elution buffer. Eluted fractions were immediately neutralized with 1 M Tris-HCl, pH 9.0 and further purified using a Superdex 200 Increase size exclusion column (GE Healthcare). Fc fusions of ACE2 and VHH-72 were purified the same way as IgGs. The VHH-72 apoferritin fusion was purified by hydrophobic interaction chromatography using a HiTrap Phenyl HP column and the eluted fraction was loaded onto a Superose 6 10/300 GL size exclusion column (GE Heathcare) in 20 mM sodium phosphate pH 8.0, 150 mM NaCl. The prefusion S ectodomain (BEI NR52394), wild type RBD (BEI NR52309) and mutant RBDs were purified using a HisTrap Ni-NTA column (GE Healthcare) followed by Superose 6 and Superdex 200 Increase size exclusion columns (GE Heathcare), respectively in 20 mM phosphate pH 8.0, 150 mM NaCl buffer.

#### Design, expression and purification of split Multabodies

Genes encoding scFab and scFc fragments linked to half apoferritin were generated by deletion of residues 1 to 95 (C-Ferritin) and 95 to 175 (N-Ferritin) of the light chain of human apoferritin using the KOD-Plus mutagenesis kit (Toyobo, Osaka, Japan). Transient transfection of the split Multabodies in HEK 293F cells were obtained by mixing 66 μg of the plasmids scFab-human apoferritin: scFc-human N-Ferritin: scFab-C-Ferritin in a 2:1:1 ratio. In the case of multispecific Multabodies, a 4:2:1:1 ratio of scFab1-human apoferritin: scFc-human N-Ferritin: scFab2-C-Ferritin: scFab3-C-Ferritin was used. The DNA mixture was filtered and incubated at RT with 66 μl of FectoPRO before adding to the cell culture. Split Multabodies were purified by affinity chromatography using a HiTrap Protein A HP column (GE Healthcare) with 20 mM Tris pH 8.0, 3 M MgCl_2_ and 10% glycerol elution buffer. Fractions containing the protein were concentrated and further purified by gel filtration on a Superose 6 10/300 GL column (GE Healthcare).

#### Negative-stain electron microscopy

3 μL of Multabody at a concentration approximately of 0.02 mg/mL was placed on the surface of a carbon-coated copper grid that had previously been glow-discharged in air for 15 sec, allowed to adsorb for 30 s, and stained with 3 μL of 2% uranyl formate. Excess stain was removed immediately from the grid using Whatman No. 1 filter paper and an additional 3 μL of 2% uranyl formate was added for 20 s. Grids were imaged with a FEI Tecnai T20 electron microscope operating at 200 kV and equipped with an Orius charge-coupled device (CCD) camera (Gatan Inc).

#### Biolayer interferometry

Direct binding kinetics measurements were conducted using an Octet RED96 BLI system (Sartorius ForteBio) in PBS pH 7.4, 0.01% BSA and 0.002% Tween at 25° C. His-tagged RBD or SARS-CoV-2 Spike protein was loaded onto Ni-NTA (NTA) biosensors (Sartorius ForteBio) to reach a BLI signal response of 0.8 nm. Association rates were measured by transferring the loaded biosensors to wells containing a 2-fold dilution series from 250 to 16 nM (Fabs), 125 to 4 nM (IgG), and 16 to 0.5 nM (MB). Dissociation rates were measured by dipping the biosensors into buffer-containing wells. The duration of each of these two steps was 180 s. Fc characterization in the split Multabody design was assessed by measuring binding to hFcγRI and hFcRn. To probe the theoretical capacity of the Multabodies to undergo endosomal recycling, binding to the hFcRn β2-microglobulin complex was measured at physiological (7.5) and endosomal (5.6) pH. Competition assays were performed in a two-step binding process. Ni-NTA biosensors preloaded with His-tagged RBD were first dipped into wells containing the primary antibody at 50 μg/mL for 180 s. After a 30 s baseline period, the sensors were dipped into wells containing the second antibody at 50 μg/ml for an additional 300 s.

#### Dynamic light scattering

The hydrodynamic radius (Rh) of the Multabody was determined by dynamic light scattering (DLS) using a DynaPro Plate Reader III (Wyatt Technology). 20 μL of the Multabody at a concentration of 1 mg/mL was added to a 384-well black, clear bottom plate (Corning) and measured at a fixed temperature of 25° C with a duration of 5 s per read. Particle size determination and polydispersity were obtained from the accumulation of 5 reads using the Dynamics software (Wyatt Technology).

#### Aggregation temperature measurements

Aggregation temperature (T_agg_) of the Multabodies and parental IgGs were determined using a UNit instrument (Unchained Labs). Samples were concentrated to 1.0 mg/mL and subjected to a thermal ramp from 25 to 95° C with 1° C increments. T_agg_ was determined as the temperature at which 50% increase in the static light scattering at a 266 nm wavelength relative to baseline was observed (i.e. the maximum value of the differential curve). The average and the standard error of two independent measurements were calculated using the UNit analysis software.

#### Pharmacokinetics and immunogenicity studies

A surrogate Multabody composed of the scFab and scFc fragments of mouse HD37 (anti-hCD19) IgG2a fused to the N-terminus of the light chain of mouse apoferritin (mFerritin) was used for the study. HD37 scFab-mFerritin: Fc-mFerritin: mFerritin in a 2:1:1 ratio was transfected and purified following the procedure described above. L234A, L235A and P329G mutations were introduced in the mouse IgG2a Fc-construct to silence effector functions of the Multabody^43^. *In vivo* studies were performed using 12-week-old male C57BL/6 mice purchased from Charles River (Strain code: 027), housed in individually-vented cages. All procedures were approved by the Local Animal Care Committee at the University of Toronto Scarborough. A single injection of approximately 5 mg/kg of Multabodies or control samples (HD37 IgG-IgG1 and IgG2a subtypes) and *Helicobacter pylori* ferritin (hpFerritin)-PfCSP malaria peptide in 200 μL of PBS (pH 7.5) were subcutaneously injected. Blood samples were collected at multiple time points and serum samples were assessed for levels of circulating antibodies and anti-drug antibodies (ADA) by ELISA. Briefly, 96-well Pierce Nickel Coated Plates (Thermo Fisher) were coated with 50 μL at 0.5 μg/ml of the His_6x_-tagged antigen hCD19 to determine circulating HD37-specific concentrations using reagent-specific standard curves for IgGs and Multabodies. For ADA determination, Nunc MaxiSorp plates (Biolegend) were coated with a 12-mer HD37 scFab-mFerritin or with the hpFerritin-PfCSP malaria peptide. HRP-ProteinA (Invitrogen) was used as a secondary molecule and the chemiluminescence signal was quantified using a Synergy Neo2 Multi-Mode Assay Microplate Reader (Biotek Instruments).

#### Biodistribution studies

8-week-old BALB/c mice were purchased from The Jackson Laboratory and housed in individually-vented caging. All procedures were approved by the Local Animal Care Committee at the University of Toronto. The HD37 IgG2a Multabody or HD37 IgG2a control were fluorescently conjugated with Alexa-647 using an Alexa Fluor™ 647 Antibody Labeling kit (Invitrogen) as per manufacturer instructions. PerkinElmer IVIS Spectrum (PerkinElmer) was used to conduct non-invasive biodistribution experiments. BALB/c mice were injected subcutaneously into the loose skin over the shoulders with approximately 5 mg/kg of the MB or control samples in 200 μL of PBS (pH 7.5) and imaged at time 0, 1h, 6h, 24h, 2d, 3d, 4d, 8d, 11d following injection. Prior to imaging, mice were placed in an anesthesia induction chamber containing a mixture of isoflurane and oxygen for 1 min. Anesthetized mice were then placed in the prone position at the center of a built-in heated docking system within the IVIS imaging system (maintained at 37°C and supplied with a mixture of isoflurane and oxygen). For whole body 2D imaging, mice were imaged for 1-2 s (excitation 640 nm, emission 680 nm) inside the imaging system. Data were analyzed and reconstructed with the IVIS software (Living Image Software for IVIS) using 640 nm/680 nm laser.

#### Panning of Phage libraries against the RBD of SARS-CoV-2

143 healthy human subjects were used for the assembly of a SuperHuman 2.0 library. CDR-H3 diversity was sourced by PCR from naive CD27-IgM+ B-cells while other CDR diversity was sourced from CD27+IgG+ B cells from the therapeutic frameworks IGHV1-46, IGHV1-69, IGHV3-15, IGHV3-23, IGKV1-39, IGKV2-28, IGKV3-15 and IGKV4-1 by using PCR overlap extension. The light and heavy chains were transformed to exceed 1e^8^ and 7.6e^10^ transformants for each chain framework, respectively. Illumina MiSeq replicate rarefaction analysis was used to assure a recovery of over 98% unique clones. In addition, Protein L and Protein A were used (10 min incubation at 70° C) to select thermostable and well-expressing light chain diversity. The EXPi-293 mammalian expression system was used for expression of the receptor Binding domain (RBD)-Fc-Avi tag construct of the SARS-CoV-2. This protein was subsequently purified by protein G Dynabeads, biotinylated and quality-controlled for biotinylation and binding to ACE2 recombinant protein (Sino Biologics Inc). The SuperHuman 2.0 Phage library (5×10^12^) was heated for 10 min at 72° C and de-selected against Protein G Dynabeads™ (Invitrogen), M-280 Streptavidin Dynabeads™ (Invitrogen), Histone from Calf Thymus (Sigma), Human IgG (Sigma) and ssDNA-Biotin NNK from Integrated DNA Technologies and DNA-Biotin NNK from Integrated DNA Technologies. Next, the library was panned against the RBD-captured by M-280 Streptavidin Dynabeads™ using an automated protocol on Kingfisher FLEX (Thermofisher). Selected phages were acid eluted from the beads and neutralized using Tris-HCl pH 7.9 (Teknova). ER2738 cells were infected with the neutralized phage pools at OD_600_=0.5 at a 1:10 ratio and after 40 min incubation at 37° C and 100 rpm, the phage pools were centrifuged and incubated on agar with antibiotic selection overnight at 30° C. The rescued phages were precipitated by PEG and subjected to three additional rounds of soluble-phase automated panning. PBST/1% BSA buffer and/or PBS/1% BSA was used in the de-selection, washes and selection rounds.

#### Screening of anti-SARS-CoV-2 scFvs in bacterial periplasmic extracts with SARS-CoV-2 RBD

Anti-SARS-CoV-2 RBD scFvs selected from phage display were expressed and screened using high-throughput surface plasmon resonance (SPR) on Carterra LSA Array SPR instrument (Carterra) equipped with HC200M sensor chip (Carterra) at 25° C. A V5 epitope tag was added to the scFv to enable capture via immobilized anti-V5 antibody (Abcam, Cambridge, MA) that was pre-immobilized on the chip surface by standard amine-coupling. Briefly: the chip surface was first activated by 10 min injection of a 1:1:1 (v/v/v) mixture of 0.4 M 1-Ethyl-3-(3-Dimethylaminopropyl) carbodiimide hydrochloride (EDC), 0.1 M N-hydroxysulfosuccinimide (sNHS) and 0.1 M 2-(N-morpholino) ethanesulfonic acid (MES) pH 5.5. Then, 50 μg/ml of anti-V5 tag antibody prepared in 10 mM sodium acetate pH 4.3 was coupled for 14 min and the excess reactive esters were blocked with 1 M ethanolamine HCl pH 8.5 during a 10 min injection. For screening, a 384-ligand array comprising of crude bacterial periplasmic extracts (PPE) containing the scFvs (1 spot per scFv) was prepared. Each extract was prepared at a 2-fold dilution in running buffer (10 mM HEPES pH 7.4, 150 mM NaCl, 3 mM EDTA, 0.01% (v/v) Tween-20 (HBSTE)) and printed on the anti-V5 surface for 15 min. SARS-CoV-2 RBD Avi Tev His tagged was then prepared at 0, 3.7, 11.1, 33.3, 100, 37, and 300 nM in 10 mM HEPES pH 7.4, 150 mM NaCl, 0.01% (v/v) Tween-20 (HBST) supplemented with 0.5 mg/ml BSA and injected as analyte for 5 min with a 15 min dissociation time. Samples were injected in ascending concentration without any regeneration step. Binding data from the local reference spots was used to subtracted signal from the active spots and the nearest buffer blank analyte responses were subtracted to double-reference the data. The double-referenced data were fitted to a simple 1:1 Langmuir binding model in Carterra’s Kinetic Inspection Tool (version Oct. 2019). 20 medium-affinity binders from phage display screening were selected for the present study.

#### Virus production and pseudovirus neutralization assays

SARS-CoV-2 pseudotyped viruses (PsV) were generated using an HIV-based lentiviral system as previously described^44^ with few modifications. Briefly, 293T cells were co-transfected with a lentiviral backbone encoding the luciferase reporter gene (BEI NR52516), a plasmid expressing the Spike (BEI NR52310) and plasmids encoding the HIV structural and regulatory proteins Tat (BEI NR52518), Gag-pol (BEI NR52517) and Rev (BEI NR52519). 24 h post transfection at 37° C, 5 mM sodium butyrate was added to the media and the cells were incubated for an additional 24-30 h at 30° C. SARS-CoV-2 Spike mutant D614G was kindly provided by D.R. Burton (The Scripps Research Institute) and the rest of the PsV mutants were generated using the KOD-Plus mutagenesis kit (Toyobo, Osaka, Japan). Neutralization was determined in a single-cycle neutralization assay using 293T-ACE2 cells (BEI NR52511) and HeLa-ACE2 cells (kindly provided by D.R. Burton; The Scripps Research Institute). PsV neutralization was monitored by adding Britelite plus reagent (PerkinElmer) to the cells and measuring luminescence in relative light units (RLUs) using a Synergy Neo2 Multi-Mode Assay Microplate Reader (Biotek Instruments). IC_50_ fold increase was calculated as: IgG_IC50_ (μg/mL) / MB_IC50_ (μg/mL). Two to three biological replicates with two technical replicates each were performed.

#### Authentic virus neutralization assays

VeroE6 cells were seeded in a 96F plate at a concentration of 30,000/well in DMEM supplemented with 100U Penicillin, 100U Streptomycin and 10% FBS. Cells were allowed to adhere to the plate and rest overnight. After 24 h, 5-fold serial dilutions of the IgG and MB samples were prepared in DMEM supplemented with 100U Penicillin and 100U Streptomycin in a 96R plate in quadruplicates (25 uL/well). 25 uL of SARS-CoV-2/SB2-P4-PB^45^ Clone 1 was added to each well at 100TCID/well and incubated for 1 h at 37 °C with shaking every 15 min. After co-culturing, the media from the VeroE6 plate was removed, and 50 uL antibody-virus sample was used to inoculate VeroE6 cells in quadruplicates for 1 h at 37 °C, 5% CO_2_, shaking every 15 min. After 1 h inoculation, the inoculum was removed and 200 uL of fresh DMEM supplemented with 100U Penicillin, 100U Streptomycin and 2% FBS was added to each well. The plates were further incubated for 5 days. The cytopathic effect (CPE) was monitored and PRISM was used to calculate IC_50_ values. Three biological replicates with four technical replicates each were performed.

#### Cross-linking of Spike protein with Fabs 80, 298 and 324

100 μg of Spike trimer was mixed with 2x molar excess of Fab 80, 298 or 324 in 20 mM HEPES pH 7.0, 150 mM NaCl. Proteins were crosslinked by addition of 0.075% (v/v) glutaraldehyde (Sigma Aldrich) and incubated at room temperature for 120 min. Complexes were purified via size exclusion chromatography (Superose6 Increase 10/300 GL, GE Healthcare), concentrated to 0.5 mg/mL and directly used for cryo-EM grid preparation.

#### Cross-linking of Fab 46-RBD complex

100 μg of Fab 46 was mixed with 2x molar excess of RBD in 20 mM HEPES pH 7.0, 150 mM NaCl. The complex was crosslinked by addition of 0.05% (v/v) glutaraldehyde (Sigma Aldrich) and incubated at room temperature for 45 min. The cross-linked complex was purified via size exclusion chromatography (Superdex 200 Increase 10/300 GL, GE Healthcare), concentrated to 2.0 mg/ml and directly used for cryo-EM grid preparation.

#### Cryo-EM data collection and image processing

3 μl of sample was deposited on holey gold grids prepared in-house^51^, which were glow-discharged in air for 15 s with a PELCO easiGlow (Ted Pella) before use. Sample was blotted for 6 s with a modified FEI Mark III Vitrobot (maintained at 4° C and 100% humidity) using an offset of −5, and subsequently plunge-frozen in a mixture of liquid ethane and propane. Data was acquired at 300 kV with a Thermo Fisher Scientific Titan Krios G3 electron microscope and prototype Falcon 4 camera operating in electron counting mode at 250 frames/s. Movies were collected for 9.6 s with 29 exposure fractions, a camera exposure rate of ~5 e^−^/pix/s, and total specimen exposure of ~44 e^−^/Å^2^. No objective aperture was used. The pixel size was calibrated at 1.03 Å/pixel from a gold diffraction standard. The microscope was automated with the *EPU* software package and data collection was monitored with *cryoSPARC Live*^52^.

To overcome preferred orientation encountered with some of the samples, tilted data collection was employed^53^. For the Spike-Fab 80 complex, 820 0° tilted movies and 2790 40° tilted movies were collected. For the Spike-Fab 298 complex, 4259 0° tilted movies and 3513 40° tilted movies were collected. For the Spike-Fab 324 complex, 1098 0° tilted movies and 3380 40° tilted movies were collected. For the RBD-Fab 46 complex, 4722 0° tilted movies were collected. For 0° tilted movies, cryoSPARC patch motion correction was performed. For 40° tilted movies, *Relion* MotionCorr^54,55^ was used. Micrographs were then imported into *cryoSPARC* and patch CTF estimation was performed. Templates generated from 2D classification during the *cryoSPARC Live* session were used for template selection of particles. 2D classification was used to remove junk particle images, resulting in a dataset of 80,951 particle images for the Spike-Fab 80 complex, 203,138 particle images for the Spike-Fab 298 complex, 64,365 particle images for the Spike-Fab 324 complex, and 2,143,629 particle images for the RBD-Fab 46 complex. Multiple rounds of multi-class *ab initio* refinement were used to clean-up the particle image stacks, and homogeneous refinement was used to obtain consensus structures. For tilted particles, particle polishing was done within *Relion* at this stage and re-imported back into *cryoSPARC*. For the Spike-Fab complexes, extensive flexibility was observed. 3D variability analysis was performed^56^ and together with heterogeneous refinement used to classify out the different states present. Non-uniform refinement was then performed on the final set of particle images^57^. For the RBD-Fab 46 complex, *cryoSPARC ab initio* refinement with three classes was used iteratively to clean up the particle image stack. Thereafter, the particle image stack with refined Euler angles was brought into *cis*TEM for reconstruction^58^ to produce a 4.0 Å resolution map. Transfer of data between *Relion* and *cryoSPARC* was done with *pyem*^59^.

#### Crystallization and structure determination

A ternary complex of 52 Fab-298 Fab-RBD was obtained by mixing 200 μg of RBD with 2X molar excess of each Fab in 20 mM TRIS pH 8.0, 150 mM NaCl and subsequently purified via size exclusion chromatography (Superdex 200 Increase 10/300 GL, GE Healthcare). Fractions containing the complex were concentrated to 7.3 mg/ml and mixed in a 1:1 ratio with 20% (w/v) 2-propanol, 20% (w/v) PEG 4000, 0.1 M sodium citrate pH 5.6. Crystals appeared after ~1 day and were cryoprotected in 10% (v/v) ethylene glycol before being flash-frozen in liquid nitrogen. Data were collected on the 23-ID-D beamline at the Argonne National Laboratory Advanced Photon Source. The dataset was processed using XDS and XPREP^46^. Phases were determined by molecular replacement using Phaser^47^ with CNTO88 Fab as a model for 52 Fab (PDB ID: 4DN3), 20358 Fab as a model for 298 Fab (PDB ID: 5CZX), and PDB ID: 6XDG as a search model for the RBD. Refinement of the structure was performed using phenix.refine^48^ and iterations of manual building and refinement in Coot^49^. Access to all software was supported through SBGrid^50^.

---

**Fig. S1.**
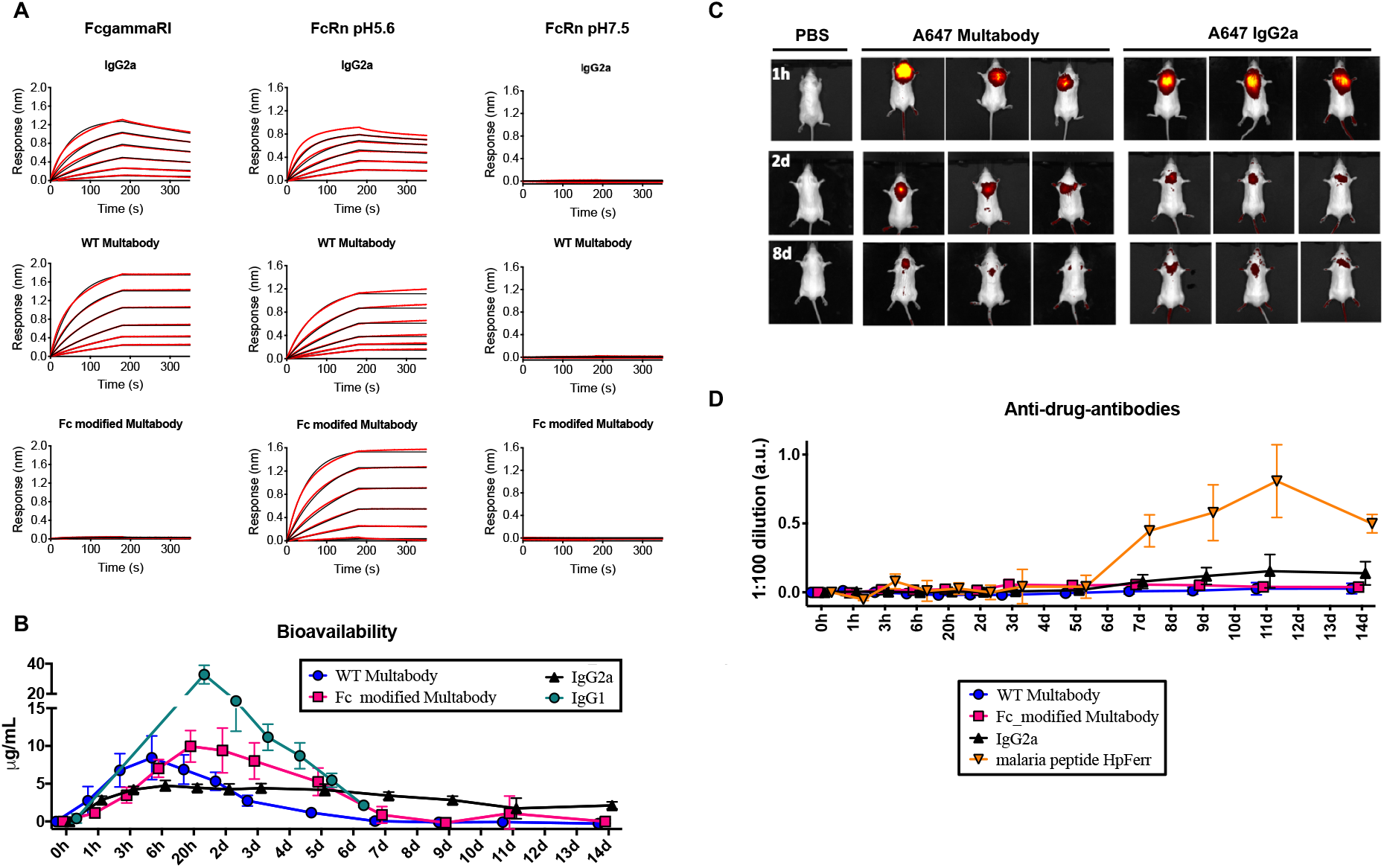
Bioavailability, biodistribution and immunogenicity of a surrogate Multabody in mice. **(A)** Binding kinetics of WT and Fc-modified (LALAP mutation) MB to mouse FcγRI (left) and mouse FcRn at endosomal (middle) and physiological (right) pH in comparison to the parental IgG. 2-fold dilution series from 100 to 3 nM (IgG), and 10 to 0.3 nM (MB) were used. Black lines represent raw data whereas red lines represent global fits. **(B)** Five male C57BL/6 mice per group were used to assess the serum concentration of a surrogate mouse MB, a Fc-modified MB (LALAP mutation) and parental mouse IgGs (IgG1 and IgG2a subtype) after subcutaneous administration of 5 mg/kg. **(C)** MB and IgG2a samples were labeled with Alexa-647 for visualization of their biodistribution post subcutaneous injection into three BALB/c mice/group via live non-invasive 2D whole body imaging. **(D)** Five male C57BL/6 mice per group were used to assess any anti-drug antibody response induced by the mouse surrogate Multabody in comparison to parental IgG and a species-mismatched malaria PfCSP peptide fused to *Helicobacter pylori* ferritin (HpFerr).

**Fig. S2.**
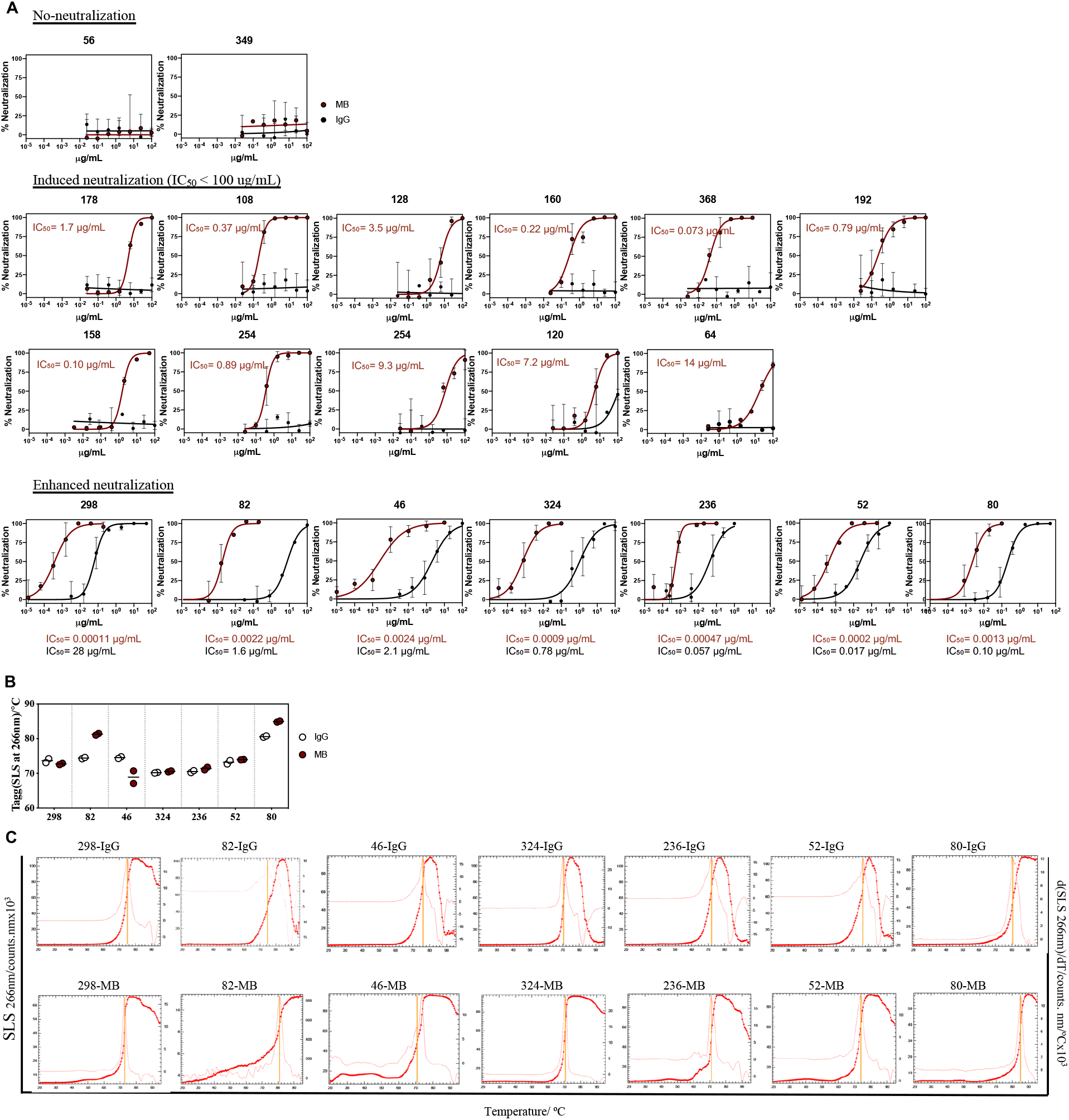
Neutralization and thermostability of SARS-CoV-2 RBD-targeting Multabodies and their parental IgGs. **(A)** Representative neutralization titration curves of 20 antibodies against SARS-CoV-2 PsV when displayed as IgGs (black) and MBs (dark red). The mean IC_50_ value of three biological replicates, each with two technical replicates are displayed for comparison. (**B**) Comparison of the aggregation temperature (T_agg_) of the seven most potent IgGs (white) and their respective MBs (dark red). (**C**) Static light scattering (SLS) at 266 nm *versus* temperature plots (dark red) from (b). T_agg_ values are calculated from the maximum of the differential curves (light red) and indicated with yellow lines.

**Fig. S3.**
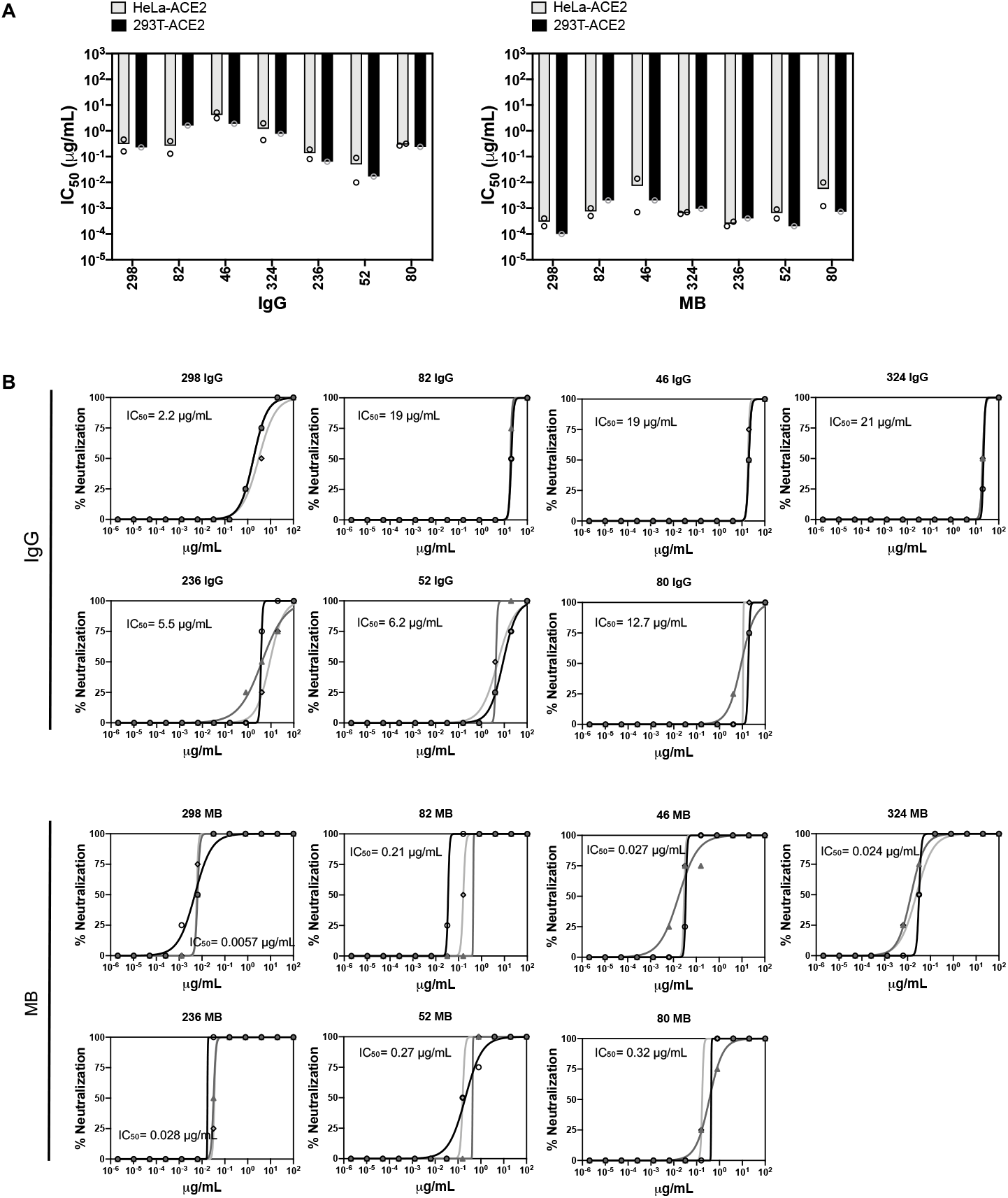
Neutralization profiles of selected IgGs and MBs in different assays. (**A**) Similar neutralization profiles of IgGs (left) vs MBs (right) against pseudotyped SARS-CoV-2 PsV targeting 293T-ACE2 (black) and HeLa-ACE2 (gray) target cells. The mean IC_50_ value and individual IC_50_ values of three and two biological replicates are shown for 293T-ACE2 and HeLa-ACE2 cells, respectively. (**B**) Neutralization titration curves of three biological replicates (different shades of gray) against the authentic SARS-CoV-2/SB2-P4-PB strain^45^. The mean IC_50_ is indicated. The less sensitive neutralization phenotype observed against authentic virus in comparison to PsV is in agreement with previous reports^5,13,14,17^. However, other studies have observed similar values^3,11,12,16^ between the two assays. This discrepancy makes cross-comparison of antibody potencies against live replicating virus difficult and is likely due to differences in the length of time of the neutralization experiment. Short incubation times will minimize the number of replications that the virus can undergo, resembling the one replication cycle of the PsV assays.

**Fig. S4.**
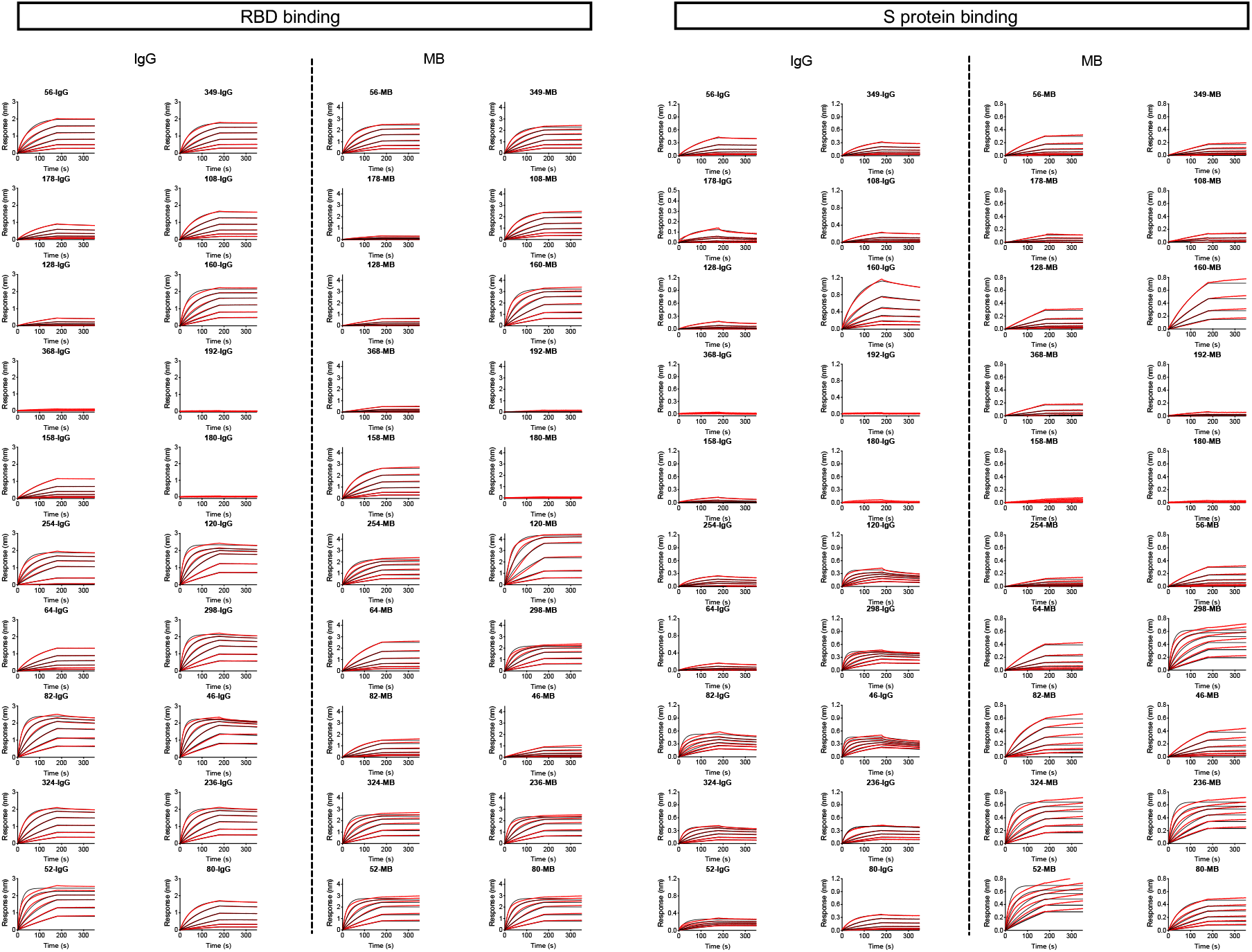
Binding profiles of IgGs and MBs. BLI response curves of IgG and MBs binding to RBD (left) and S protein (right) of SARS-CoV-2 immobilized onto Ni-NTA biosensors. 2-fold dilution series from 125 to 4 nM (IgG), and 16 to 0.5 nM (MB) were used. Black lines represent raw data whereas red lines represent global fits.

**Fig. S5.**
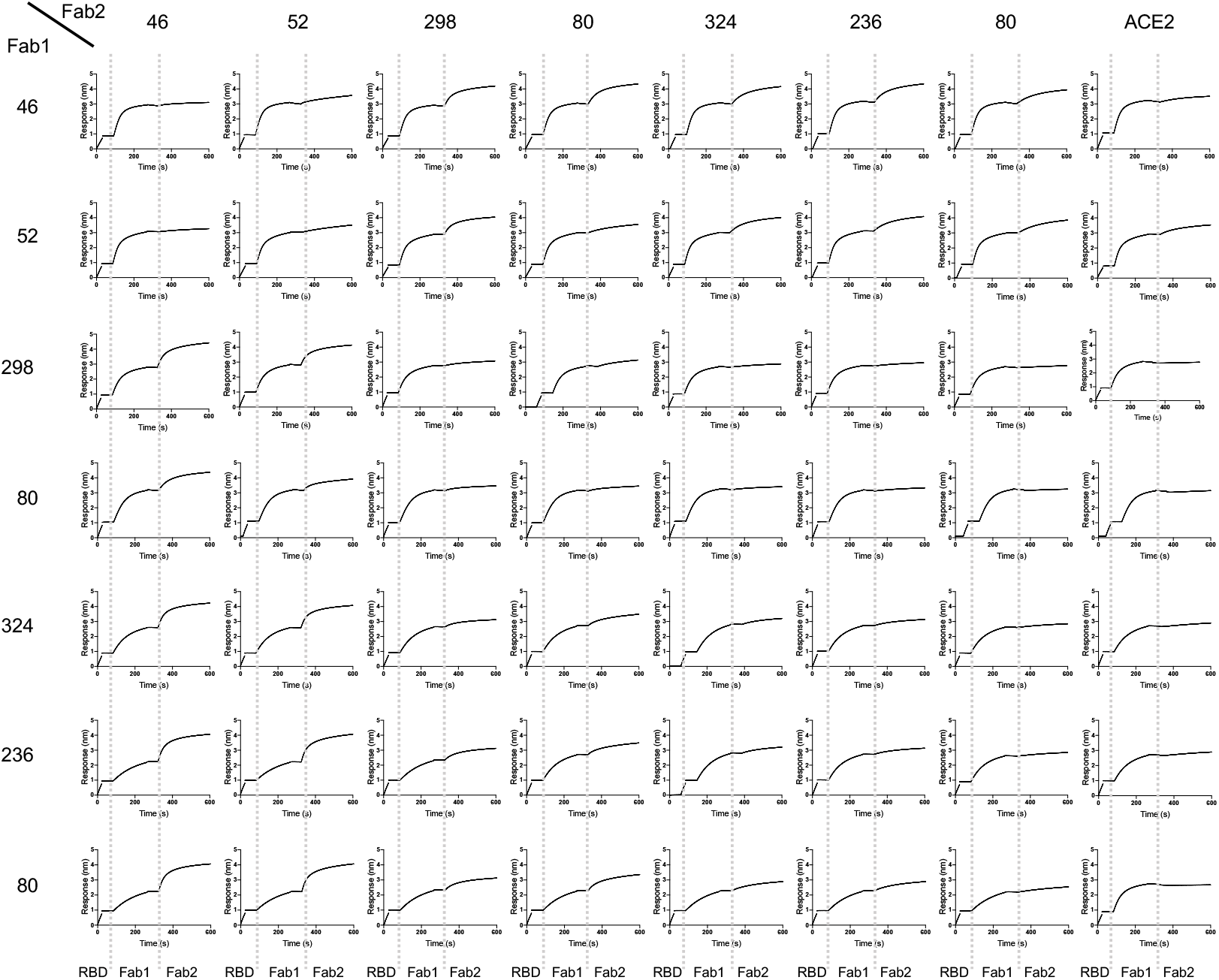
Epitope binning. mAb binding competition experiments to His-tagged RBD as measured by biolayer interferometry (BLI). 50 μg/ml of mAb 1 was incubated for 3 min followed by incubation with 50 μg/ml of mAb 2 for 5 min.

**Fig. S6.**
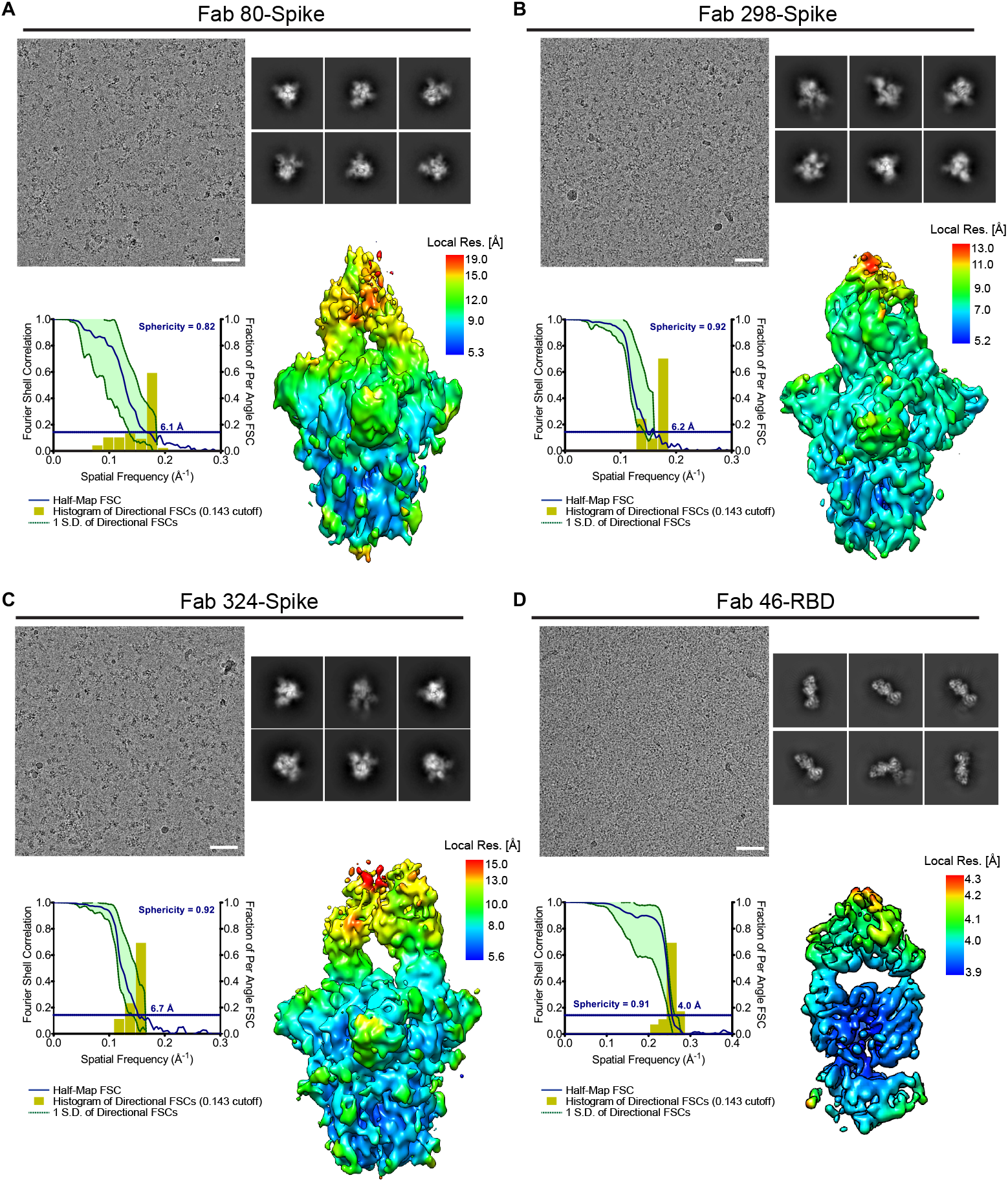
Cryo-EM analysis of the Fab-Spike and Fab-RBD complexes. Representative cryo-EM micrograph (Scale bar 50 nm, top left), selected 2D class averages (top right), Fourier shell correlation curve from the final 3D non-uniform refinement (bottom left) and local resolution (Å) plotted on the surface of the cryo-EM map (bottom right) are shown for the Fab 80-Spike complex (**A**), the Fab 298-Spike complex (**B**), the Fab 324-Spike complex (**C**), and the Fab 46-RBD complex (**D**).

**Fig. S7.**
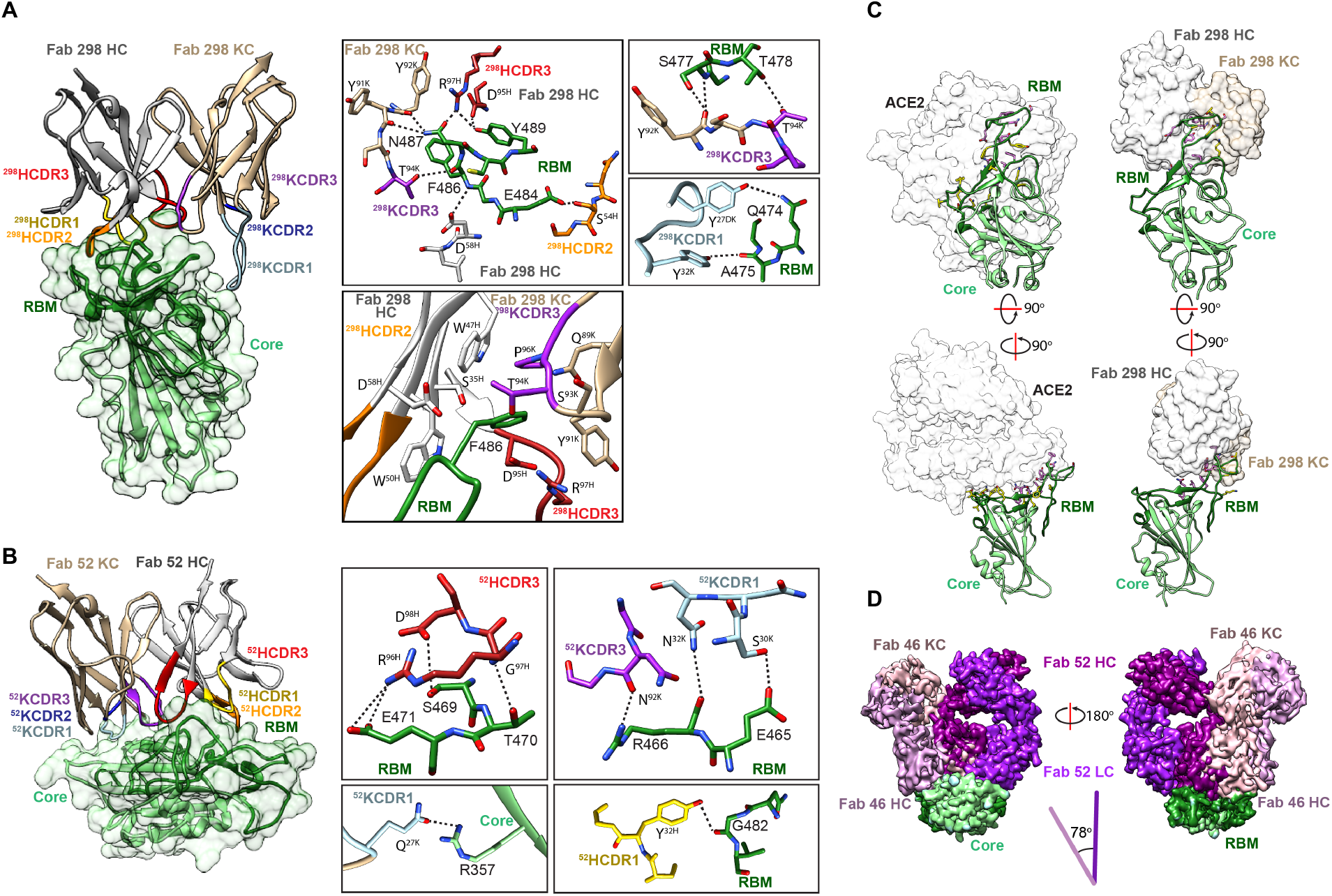
Binding interfaces of mAbs 52 and 298 and the RBD. Interaction of mAbs 298 (**A**) and 52 (**B**) with RBD (light and dark green for the core and RBM regions, respectively) is mediated by complementarity determining regions (CDR) heavy (H) 1 (yellow), H2 (orange), H3 (red), kappa light (K) 1 (light blue) and K3 (purple) (left panels). Critical binding residues are shown in sticks (right panels). H-bonds and salt bridges are depicted as black dashed lines. L and H chains of Fabs are shown in tan and white, respectively. (**C**) Bottom and side views of ACE2 (left) and Fab 298 (right) bound to RBD. RBD side-chains that are part of the binding interface of the ACE2-RBD and Fab 298-RBD complexes are depicted in pink, while RBD side-chains unique to a given interface are shown in yellow. Surfaces of ACE2, variable regions of Fab 298 HC and Fab 298 KC are shown in white, grey and tan, respectively. The RBD is colored as in (a). (**D**) Superposition of Fabs 46 (light pink) and 52 (dark pink) when bound to the RBD (green) reveals a distinct angle of approach for the two mAbs.

**Fig. S8.**
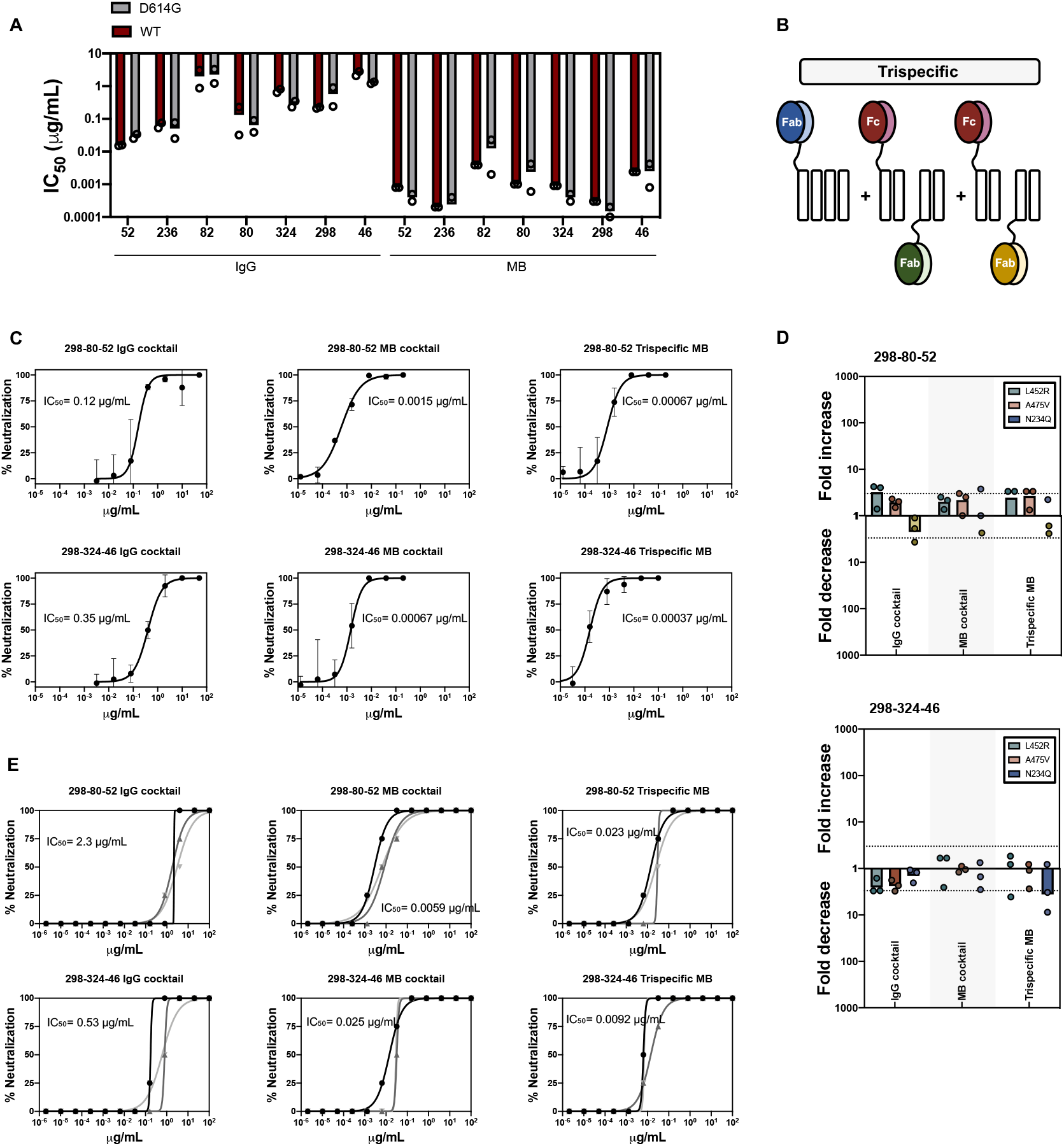
The MB platform potently overcomes SARS-CoV-2 sequence variability. **(A)** Comparison of the neutralization potency of selected IgGs and MBs against WT PsV (dark red) and the more infectious D614G PsV (grey). (**B**) Schematic representation of a tri-specific MB generated by combination of three Fab specificities and the Fc fragment using the MB split design. (**C**) Cocktails and tri-specific MBs that combine the specificities of mAbs 298, 80 and 52, or 298, 324 and 46 were generated and tested against WT PsV. (**D**) Neutralization potency change of cocktails and tri-specific MBs against pseudotyped SARS-CoV-2 variants in comparison to WT PsV. PsV variants that were sensitive to individual antibodies within the cocktails were selected. The area within the dotted lines represent a 3-fold change in IC_50_ value. This threshold was established as the cutoff to establish increased sensitivity (up bars) and increased resistance (down bars). (**E**) Neutralization titration curves showing three biological replicates of cocktails and tri-specific MBs against the authentic SARS-CoV-2/SB2-P4-PB strain^45^. Mean IC_50_ values of three biological replicates are shown.

**Table S2.**
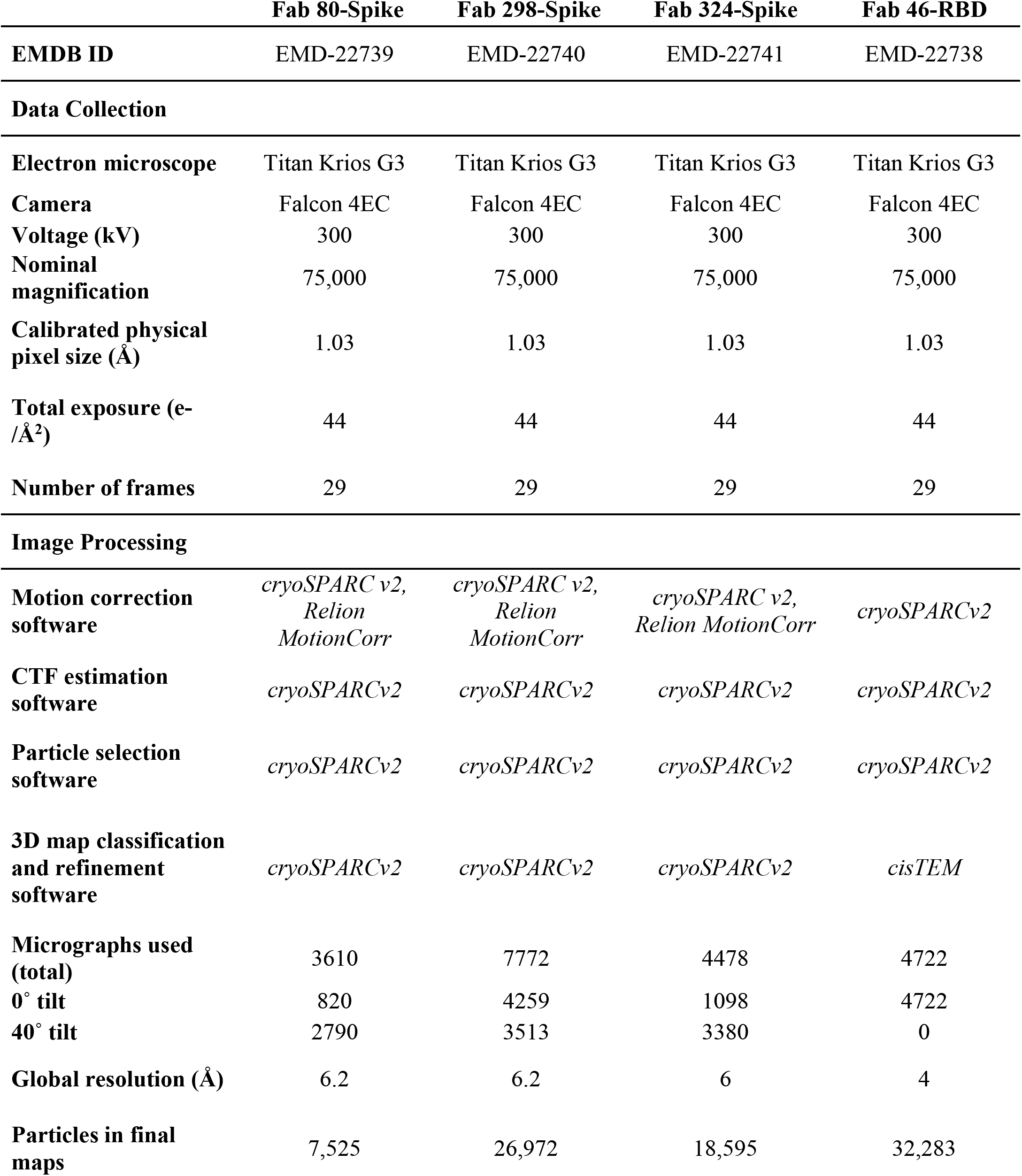
Cryo-EM data collection and image processing

**Table S3.** X-ray crystallography data collection and refinement statistics

|  | **Fab 52- Fab 298-**<br>**SARS-CoV-2 RBD** |
| --- | --- |
| **PDB ID** | 7K9Z |
| **Data Collection** |  |
| Wavelength (Å) | 1.03317 |
| Space group | P 3<sub>2</sub> 2 1 |
| Cell dimensions |  |
| a,b,c (Å) | 87.6, 87.6, 325.1 |
| $\alpha, \beta, \gamma$ (°) | 90.0, 90.0, 120.0 |
| Resolution (Å) | 39.66-2.95 (3.05-2.95) |
| No. Molecules in ASU | 1 |
| No. Total observations | 496,550 (43,958) |
| No unique observations | 31,545 (3,060) |
| Multiplicity | 15.7 (14.3) |
| R<sub>merge</sub> (%) | 16.8 (74.2) |
| R<sub>pim</sub> (%) | 4.3 (20.1) |
| <I/σI> | 12.3 (1.4) |
| CC<sub>1/2</sub> | 99.8 (86.3) |
| Completeness (%) | 99.9 (99.9) |
| **Refinement** |  |
| Non-hydrogen atoms | 8061 |
| Macromolecule | 8047 |
| Glycan | 14 |
| R<sub>work</sub>/R<sub>free</sub> | 0.259/0.286 |
| Rms deviations |  |
| Bond lengths (Å) | 0.002 |
| Bond angles (°) | 0.53 |
| Ramachandran plot |  |
| Favored regions (%) | 95.6 |
| Allowed regions (%) | 4.1 |
| Outliers (%) | 0.3 |
| Rotamer Outliers (%) | 2.6 |
| B-factors (Å<sup>2</sup>) |  |
| Wilson B-factor | 78.6 |
| Average B-factors | 103.9 |
| Average macromolecule | 103.9 |
| Average glycan | 114.3 |

**Table S4.**
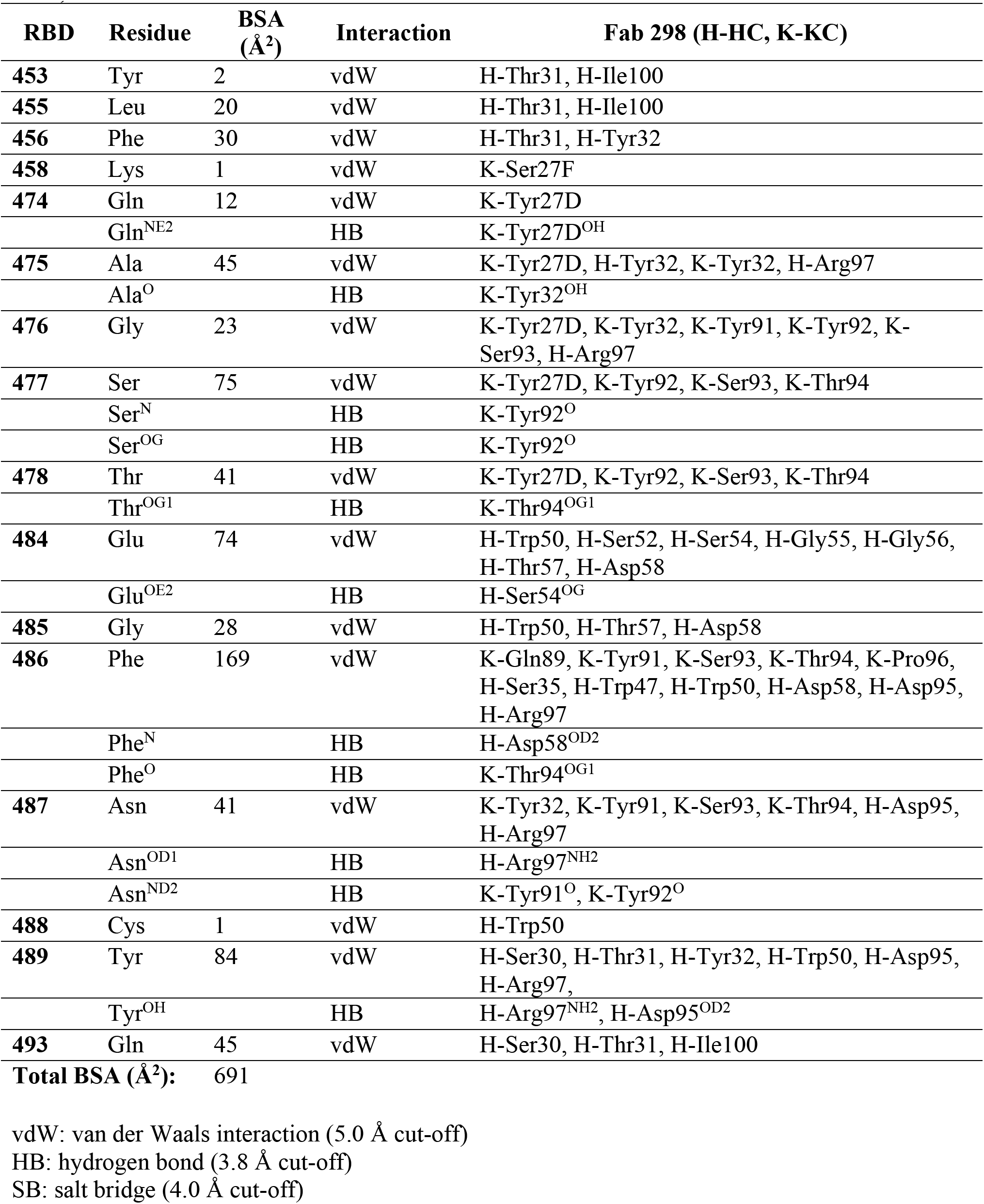
RBD residues contacting Fab 298 identified by PISA (Krissinel and Henrick, 2007).

**Table S5.**
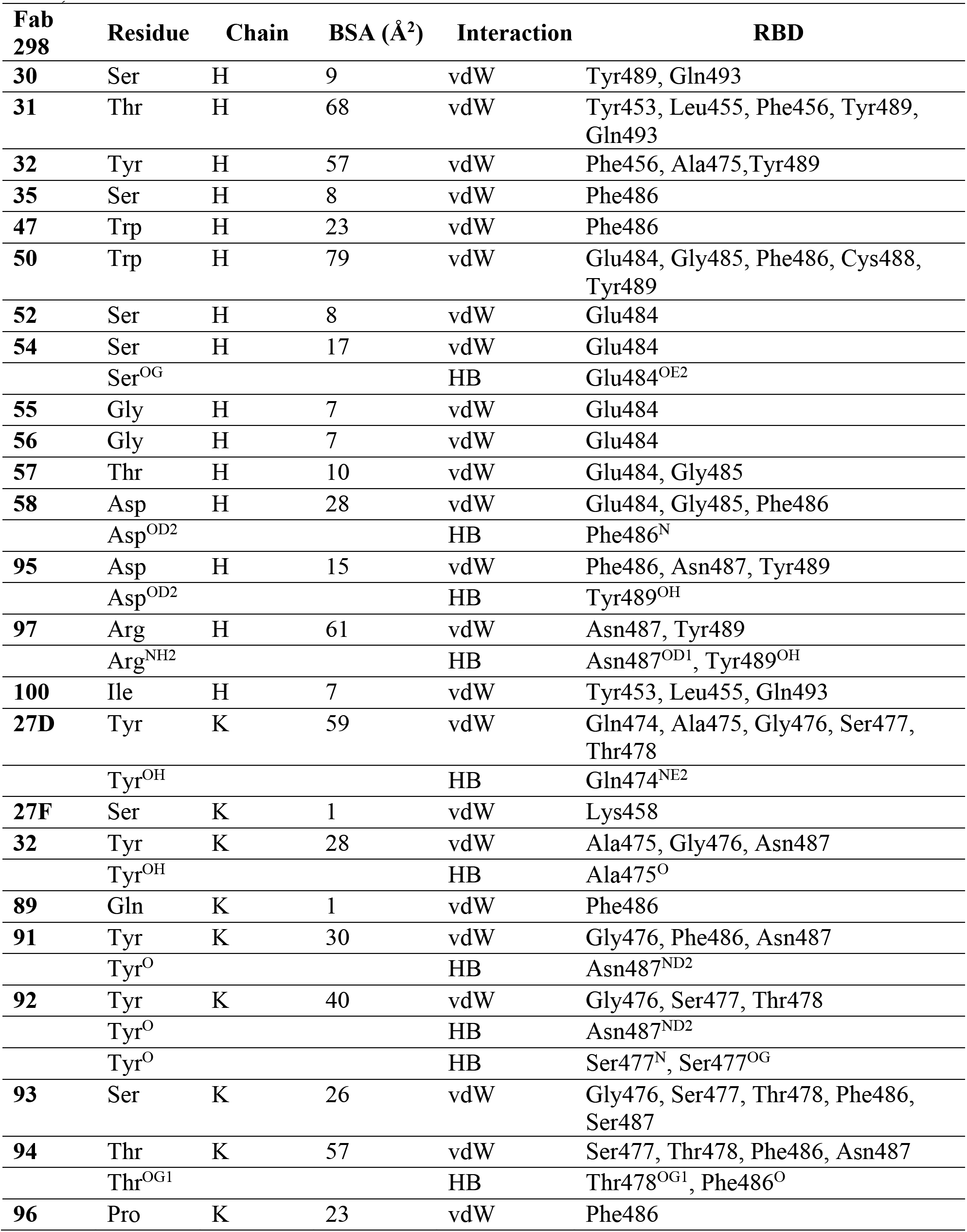
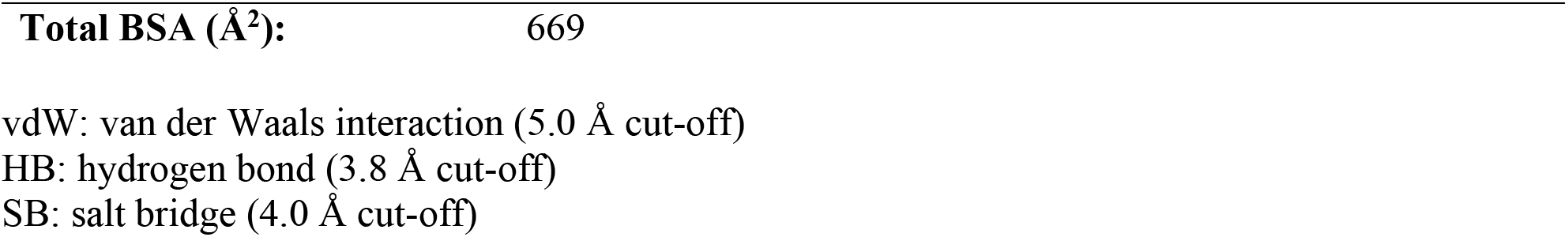
Fab 298 residues contacting RBD identified by PISA (Krissinel and Henrick, 2007).

| **Fab 298** | **Residue** | **Chain** | **BSA (Å<sup>2</sup>)** | **Interaction** | **RBD** |
| --- | --- | --- | --- | --- | --- |
| **30** | Ser | H | 9 | vdW | Tyr489, Gln493 |
| **31** | Thr | H | 68 | vdW | Tyr453, Leu455, Phe456, Tyr489, Gln493 |
| **32** | Tyr | H | 57 | vdW | Phe456, Ala475, Tyr489 |
| **35** | Ser | H | 8 | vdW | Phe486 |
| **47** | Trp | H | 23 | vdW | Phe486 |
| **50** | Trp | H | 79 | vdW | Glu484, Gly485, Phe486, Cys488, Tyr489 |
| **52** | Ser | H | 8 | vdW | Glu484 |
| **54** | Ser | H | 17 | vdW | Glu484 |
|  | Ser<sup>OG</sup> |  |  | HB | Glu484<sup>OE2</sup> |
| **55** | Gly | H | 7 | vdW | Glu484 |
| **56** | Gly | H | 7 | vdW | Glu484 |
| **57** | Thr | H | 10 | vdW | Glu484, Gly485 |
| **58** | Asp | H | 28 | vdW | Glu484, Gly485, Phe486 |
|  | Asp<sup>OD2</sup> |  |  | HB | Phe486<sup>N</sup> |
| **95** | Asp | H | 15 | vdW | Phe486, Asn487, Tyr489 |
|  | Asp<sup>OD2</sup> |  |  | HB | Tyr489<sup>OH</sup> |
| **97** | Arg | H | 61 | vdW | Asn487, Tyr489 |
|  | Arg<sup>NH2</sup> |  |  | HB | Asn487<sup>OD1</sup>, Tyr489<sup>OH</sup> |
| **100** | Ile | H | 7 | vdW | Tyr453, Leu455, Gln493 |
| **27D** | Tyr | K | 59 | vdW | Gln474, Ala475, Gly476, Ser477, Thr478 |
|  | Tyr<sup>OH</sup> |  |  | HB | Gln474<sup>NE2</sup> |
| **27F** | Ser | K | 1 | vdW | Lys458 |
| **32** | Tyr | K | 28 | vdW | Ala475, Gly476, Asn487 |
|  | Tyr<sup>OH</sup> |  |  | HB | Ala475<sup>O</sup> |
| **89** | Gln | K | 1 | vdW | Phe486 |
| **91** | Tyr | K | 30 | vdW | Gly476, Phe486, Asn487 |
|  | Tyr<sup>O</sup> |  |  | HB | Asn487<sup>ND2</sup> |
| **92** | Tyr | K | 40 | vdW | Gly476, Ser477, Thr478 |
|  | Tyr<sup>O</sup> |  |  | HB | Asn487<sup>ND2</sup> |
|  | Tyr<sup>O</sup> |  |  | HB | Ser477<sup>N</sup>, Ser477<sup>OG</sup> |
| **93** | Ser | K | 26 | vdW | Gly476, Ser477, Thr478, Phe486, Ser487 |
| **94** | Thr | K | 57 | vdW | Ser477, Thr478, Phe486, Asn487 |
|  | Thr<sup>OG1</sup> |  |  | HB | Thr478<sup>OG1</sup>, Phe486<sup>O</sup> |
| **96** | Pro | K | 23 | vdW | Phe486 |

|  |  |
| --- | --- |
| **Total BSA ($\text{\AA}^2$):** | 669 |

vdW: van der Waals interaction (5.0 $\text{\AA}$ cut-off)
HB: hydrogen bond (3.8 $\text{\AA}$ cut-off)
SB: salt bridge (4.0 $\text{\AA}$ cut-off)

**Table S6.** RBD residues contacting Fab 52 identified by PISA (Krissinel and Henrick, 2007).

| RBD | Residue | BSA<br>(Å<sup>2</sup>) | Interaction | Fab 52 (H-HC, K-KC) |
| --- | --- | --- | --- | --- |
| 346 | Arg | 45 | vdW | H-Gln64 |
| 351 | Tyr | 29 | vdW | K-Phe94, H-Ile52, H-Thr56, H-Asn58 |
| 352 | Ala | 19 | vdW | K-Gly93, K-Phe94 |
| 354 | Asn | 6 | vdW | K-Phe94 |
| 355 | Arg | 36 | vdW | K-Ser0, K-Gln27 |
|  | Arg<sup>O</sup> |  | HB | K-Ser0<sup>OG</sup> |
| 356 | Lys | 13 | vdW | K-Ser0 |
| 357 | Arg | 66 | vdW | K-Ser0, K-Gln27 |
|  | Arg<sup>N</sup> |  | HB | K-Ser0<sup>OG</sup> |
|  | Arg<sup>NH2</sup> |  | HB | K-Gln27<sup>OE1</sup> |
| 449 | Tyr | 13 | vdW | H-Phe554, H-Thr56 |
| 450 | Asn | 38 | vdW | H-Phe554, H-Thr56 |
| 452 | Leu | 47 | vdW | H-Ile52, H-Phe554, H-Gly55, H-Thr56 |
| 462 | Lys | 1 | vdW | K-Ser30 |
| 465 | Glu | 28 | vdW | K-Ser30, K-Asn31, K-Asn32, K-Asn92 |
|  | Glu<sup>OE2</sup> |  | HB | K-Ser30<sup>OG</sup> |
| 466 | Arg | 70 | vdW | K-Asn32, K-Asn92, K-93Gly, K-Phe94 |
|  | Arg<sup>O</sup> |  | HB | K-Asn32<sup>ND2</sup> |
|  | Arg<sup>NH1</sup> |  | HB | K-Asn92<sup>O</sup> |
| 467 | Asp | 12 | vdW | K-Asn32, K-Asn92, H-Asp98 |
| 468 | Ile | 102 | vdW | K-Asn32, K-Gly91, K-Asn92, K-93Gly, K-Phe94, K-Leu96, H-Arg96, H-Gly97, H-Asp98 |
| 469 | Ser | 37 | vdW | K-Asn32, H-Arg96, H-Gly97, H-Asp98 |
|  | Ser<sup>OG</sup> |  | HB | H-Asp98<sup>OD1</sup> |
| 470 | Thr | 67 | vdW | H-Ser31, H-Tyr32, H-Gly33, H-Ile52, H-Asp95, H-Arg96, H-Gly97, H-Asp98 |
|  | Thr<sup>OG1</sup> |  | HB | H-Gly97<sup>N</sup> |
| 471 | Glu | 39 | vdW | H-Tyr32, H-Arg96, H-Gly97, H-Asp98 |
|  | Glu<sup>OE1</sup> |  | SB | H-Arg96<sup>NH2</sup> |
|  | Glu<sup>OE2</sup> |  | SB | H-Arg96<sup>NH2</sup> |
| 472 | Ile | 7 | vdW | H-Ser31, H-Tyr32, H-Arg96 |
| 481 | Asn | 1 | vdW | H-Tyr32 |
| 482 | Gly | 42 | vdW | H-Ser31, H-Tyr32 |
|  | Gly<sup>O</sup> |  | HB | H-Tyr32<sup>OH</sup> |
| 483 | Val | 12 | vdW | H-Thr28, H-Ser31, H-Tyr32 |
| 484 | Glu | 53 | vdW | H-Thr28, H-Phe29, H-Thr30, H-Ser31, H-Met54 |
| 490 | Phe | 87 | vdW | H-Thr30, H-Ser31, H-Tyr32, H-Gly33, H-Ile52, H-Met54, H-Phe554 |
| 492 | Leu | 13 | vdW | H-Ile52, H-Met54, H-Phe554 |
| 493 | Gln | 8 | vdW | H-Phe554 |
| **494** | Ser | 17 | vdW | H-Phe554 |
| **Total BSA (Å<sup>2</sup>):** |  | 904 |  |  |

vdW: van der Waals interaction (5.0 Å cut-off)
HB: hydrogen bond (3.8 Å cut-off)
SB: salt bridge (4.0 Å cut-off)

**Table S7.** Fab 52 residues contacting RBD identified by PISA (Krissinel and Henrick, 2007).

| **Fab 52** | **Residue** | **Chain** | **BSA (Å<sup>2</sup>)** | **Interaction** | **RBD** |
| --- | --- | --- | --- | --- | --- |
| **28** | Thr | H | 24 | vdW | Val483, Glu484 |
| **30** | Thr | H | 3 | vdW | Glu484, Phe490 |
| **31** | Ser | H | 66 | vdW | Glu471, Gly482, Val483, Glu484, Phe490 |
| **32** | Tyr | H | 51 | vdW | Thr470, Glu471, Ile472, Asn481, Gly482, Val483, Phe490 |
|  | Tyr<sup>OH</sup> | H |  | HB | Gly482<sup>O</sup> |
| **33** | Gly | H | 7 | vdW | Thr470, Phe490 |
| **52** | Ile | H | 62 | vdW | Tyr351, Leu452, Thr470, Phe490, Leu492 |
| **53** | Met | H | 38 | vdW | Glu484, Phe490, Leu492 |
| **54** | Phe | H | 110 | vdW | Tyr449, Asn450, Leu452, Phe490, Leu492, Gln493, Ser494 |
| **55** | Gly | H | 4 | vdW | Leu452 |
| **56** | Thr | H | 57 | vdW | Tyr351, Tyr449, Asn450, Leu452 |
| **58** | Asn | H | 10 | vdW | Tyr351 |
| **64** | Gln | H | 43 | vdW | Arg346 |
| **95** | Asp | H | 2 | vdW | Thr470 |
| **96** | Arg | H | 64 | vdW | Ile468, Ser469, Thr470, Glu471, Ile472 |
|  | Arg<sup>NH2</sup> | H |  | SB | Glu471<sup>OE1</sup>, Glu471<sup>OE2</sup> |
| **97** | Gly | H | 64 | vdW | Ile468, Ser469, Thr470, Glu471 |
|  | Gly<sup>N</sup> |  |  | HB | Thr470<sup>OG1</sup> |
| **98** | Asp | H | 28 | vdW | Asp467, Ile468, Ser469, Thr470, Glu471 |
|  | Asp<sup>OD1</sup> |  |  | HB | Ser469<sup>OG</sup> |
| **0** | Ser | K | 77 | vdW | Arg355, Lys356, Arg357 |
|  | Ser<sup>OG</sup> |  |  | HB | Arg355<sup>O</sup>, Arg357<sup>N</sup> |
| **27** | Gln | K | 60 | vdW | Arg355, Arg357 |
|  | Gln<sup>OE1</sup> |  |  | HB | Arg357<sup>NH2</sup> |
| **30** | Ser | K | 28 | vdW | Lys462, Glu465 |
|  | Ser<sup>OG</sup> |  |  | HB | Glu465<sup>OE2</sup> |
| **32** | Asn | K | 19 | vdW | Glu465, Arg466, Asp467, Ile468, Ser469 |
|  | Asn<sup>ND2</sup> |  |  | HB | Arg466<sup>O</sup> |
| **91** | Gly | K | 18 | vdW | Ile468 |
| **92** | Asn | K | 56 | vdW | Glu465, Arg466, Asp467, Ile468 |
|  | Asn<sup>O</sup> |  |  | HB | Arg466<sup>NH1</sup> |
| **93** | Gly | K | 22 | vdW | Ala352, Arg466, Ile468 |
| **94** | Phe | K | 56 | vdW | Tyr351, Ala352, Asn354, Arg466, Ile468 |
| **96** | Leu | K | 5 | vdW | Ile468 |
| **Total BSA (Å<sup>2</sup>):** |  |  | 974 |  |  |

vdW: van der Waals interaction (5.0 Å cut-off)
HB: hydrogen bond (3.8 Å cut-off)
SB: salt bridge (4.0 Å cut-off)

